# A penultimate classification of canonical antibody CDR conformations

**DOI:** 10.1101/2022.10.12.511988

**Authors:** Simon Kelow, Bulat Faezov, Qifang Xu, Mitchell Parker, Jared Adolf-Bryfogle, Roland L. Dunbrack

## Abstract

Antibody complementarity determining regions (CDRs) are loops within antibodies responsible for engaging antigens during the immune response and in antibody therapeutics and laboratory reagents. Since the 1980s, the conformations of the hypervariable CDRs have been structurally classified into a number of “canonical conformations” by Chothia, Lesk, Thornton, and others. In 2011 (North et al, J Mol Biol. 2011), we produced a quantitative clustering of approximately 300 structures of each CDR based on their length, a dihedral angle metric, and an affinity propagation algorithm. The data have been made available on our PyIgClassify website since 2015 and have been widely used in assigning conformational labels to antibodies in new structures and in molecular dynamics simulations. In the years since, it is has become apparent that many of the clusters are not “canonical” since they have not grown in size and still contain few sequences. Some clusters represent multiple conformations, given the assignment method we have used since 2015. Electron density calculations indicate that some clusters are due to misfitting of coordinates to electron density. In this work, we have performed a new statistical clustering of antibody CDR conformations. We used Electron Density in Atoms (EDIA, Meyder et al., 2017) to produce data sets with different levels of electron density validation. Clusters were chosen by their presence in high electron density cutoff data sets and with sufficient sequences (≥10) across the entire PDB (no EDIA cutoff). About half of the North et al. clusters have been “retired” and 13 new clusters have been identified. We also include clustering of the H4 and L4 CDRs, otherwise known as the “DE loop” which connects strands D and E of the variable domain. The DE loop sometimes contacts antigens and affects the structure of neighboring CDR1 and CDR2 loops. The current database contains 6,486 PDB antibody entries. The new clustering will be useful in the analysis and development of new antibody structure prediction and design algorithms based on rapidly emerging techniques in deep learning. The new clustering data are available at http://dunbrack2.fccc.edu/PyIgClassify2.

## Introduction

Antibodies are integral molecules in the process of immunity, and have also found important use as reagents in molecular biology research. Antibodies are multi-domain globular protein structures that contain constant domains that interact with immune effector cells to incite immune response to various antigens (Williams and Barclay 1988, Harpaz and Chothia 1994), and variable domains with a V-type immunoglobulin protein fold (Bork et al. 1994), which is the structural element responsible for binding antigens in human blood serum and tissue. Typically antibody antigen binding sites consist of dimers, where the antibody light chain is paired with the antibody heavy chain, but cases exists of heavy chain monomers from camelids (Hamers-Casterman et al. 1993, Arbabi Ghahroudi et al. 1997), as well as light chain homodimers (Bence-Jones antibodies) (Wu and Kabat 1970). Antibodies undergo a process called V(D)J recombination to form their genetic diversity (Dildrop et al. 1982, Tonegawa 1983). In B-cells, the variable (V), diversity (D), and joining (J) gene come together to form the binding region of the heavy chain. On the light chain, only V and J regions rearrange to form the binding region. Antibodies that successfully bind antigen and elicit an immune response are selected for in B-cells, and undergo a process called somatic mutation (Tonegawa 1983), selecting for antibodies which bind antigen with higher affinity.

The typical antigen binding site on antibodies consists of six loops, three from each variable domain, called the complementarity determining regions or CDRs (Wu and Kabat 1970). Two additional loops, adjacent to CDR1 and CDR2, which connect the ‘d’ and ‘e’ strands (and so called the “de loop” or “CDR4”), sometimes come in contact with the antigen, especially in the presence of somatic insertions in the de loop, predominantly in HIV gp120 antibodies (Kelow et al. 2020). Further, we have shown that they affect the structures and cluster choices of CDR1 and/or CDR2 (Kelow et al. 2020). CDRs 1, 2, and 4 are encoded by the V-region gene segments while CDR3 is the product of VDJ or VJ recombination in heavy chain or light chains respectively.

The canonical six CDRs were first identified by the hypervariable nature of their sequences compared to the rest of the variable domain (Kabat and Wu 1971). The first solved antibody structures began to shed light about the structural form of the CDRs (Amit et al. 1986, Sheriff et al. 1987). Near the end of the 1980s, research primarily from Cyrus Chothia and Arthur Lesk deepened our understanding of the structures of the CDRs, and in turn how CDR structure affects antibody-antigen binding. This work culminated in the understanding that CDRs take on ‘canonical’ conformations (Chothia and Lesk 1987, Chothia et al. 1989, Al-Lazikani et al. 1997), or frequently observed conformations of the CDR backbone, and gave a categorization of these canonical conformations for each CDR. In this observation, Chothia et al. established the first structural bioinformatics analysis of the hypervariable region. Even with a limited amount of structures they were able to establish the idea that even amongst a diverse number of sequences, the conformational landscape of CDR backbones are discrete enough to predict structure from sequence by choosing from a set of typically observed conformations with specific residue types at certain positions.

There have been additional research studies aimed at providing classifications of the antibody CDRs (Martin and Thornton 1996, Shirai et al. 1996, Oliva et al. 1998, Whitelegg and Rees 2004, North et al. 2011, Dunbar et al. 2014, Nikoloudis et al. 2014, Adolf-Bryfogle et al. 2015, Nowak et al. 2016). Whereas Chothia had dozen of antibodies to observe, as of October 2022 the PDB contains approximately 6,500 entries containing antibodies. As the number of structures has increased, clustering methods have grown more quantitative and sophisticated. In 1996, Martin and Thornton provided an algorithmic method using a least-squares clustering method in dihedral space and a subsequent clustering in root-mean-square deviation (RMSD) space using the Cartesian coordinates of the atoms that define the backbone of the CDR (Martin and Thornton 1996). This work provided an updated classification of the CDR clusters, but also introduced the idea of automation to the CDR classification problem that would prove useful as the number of antibody structures continued to rise.

In 2011, we used an internal dihedral angle clustering metric combined with an affinity propagation clustering algorithm (North et al. 2011). We defined the CDRs taking into account structural variation of Cα atom positions after superposition of light-chain and heavy-chain variable domains. We defined the CDRs boundaries to be the same positions in light and heavy chain domains (with the exception of the C-terminal end of L2, which is 3 residues longer than H2 to account for some structural variation following L2). After defining the clusters and cluster centroids for each length of each CDR available in the PDB, we used a cutoff of 40° for the average dihedral angle difference from the centroid of each cluster to assign CDRs in all antibody PDB structures to each of these structural families. The average is calculated over ϕ and ψ for the same CDR of the same length for the same cis-trans pattern, e.g. for L3 of length 9 with a cis residue at position 7. Our nomenclature is simple, consisting of the CDR, its length, and a cluster number based on the size of the cluster (1,2,3, etc. from largest to smallest, at the time of clustering in 2011). For example, L1 length clusters were named L1-11-1, L1-11-2, and L1-11-3. Clusters with cis peptide bonds identified the cis peptide bond explicitly, e.g. L3-9-cis7-1. Our antibody conformational clusters have been made available on our PyIgClassify database website (Adolf-Bryfogle et al. 2015), which was last updated in late 2019.

In 2016, Nowak et al. used a length-independent Cartesian RMSD metric alongside hierarchical clustering and Density-based spatial clustering of applications with noise (DBSCAN) (Ester et al., 1996) to establish canonical families of length-independent CDR structures (Nowak et al. 2016). In 2020, we applied a dihedral metric, electron density validation, and DBSCAN to cluster the conformations of CDR4 in the heavy and light chains (“H4” and “L4”) (Kelow et al. 2020). For the standard length 6 L4 loop, we found four clusters – two consisting of only L4 loops from kappa domains, one from only lambda chains, and one mixed kappa/lambda cluster. For H4, almost all structures in the PDB have a length 8 conformation from a single cluster (“H4-8-1”). A small number of germlines have other CDR4 lengths, and we defined clusters H4-6-1, H4-7-1, and L4-8-1. L4-8-1 is identical in conformation to H4-8-1.

Many antibody computational design programs have been developed and released (Baran et al. 2017, Adolf-Bryfogle et al. 2018, Chowdhury et al. 2018). Our program, RosettaAntibodyDesign (Adolf-Bryfogle et al. 2018), uses our clusters to sample CDR conformations and sequences for affinity maturation of existing antibodies, taking advantage of the sequence and structural variation observed within each of our clusters in our PyIgClassify database (Adolf-Bryfogle et al. 2015). Deep learning has overtaken other methods for protein structure prediction (Jumper et al. 2021) and design (Ovchinnikov and Huang 2021) and recently been applied to antibody structure prediction (Ruffolo et al. 2021, Lee et al. 2022) and design (Mason et al. 2021). A contemporary and rigorous understanding of the sequence-structure relationships of antibody CDRs from experimental structures will be of value in understanding the evolution of antibody specificity and in the development and interpretation of deep learning approaches to antibody structure prediction and design.

The necessity to revisit clustering of CDR conformations has become evident in recent years. Since our clustering in 2011, the number of antibody structures in the PDB has grown more than sevenfold. In 2011, we implemented B-factor (a generous value of 80), resolution (2.8 Å), and conformational energy cutoffs to filter an initial set of ~1300 structures of each CDR down to about 300 non-redundant structures for each CDR as input to clustering. However, out of the 72 non-H3 clusters that we defined, 21 of them still contain fewer than 10 unique sequences, bringing doubt on whether they should be termed “canonical clusters.” Also, with a cutoff of 40° for the average dihedral angle difference from the median structures, 11 of our clusters contain a majority of structures with average dihedral differences of greater than 30° from the median, indicating either a poor choice for the median, the mixture of two or more conformational states, or data inconsistent with the identification of clusters. Structures at 30° or more away from the centroid usually look visually very different from the centroid. The sequence-structure correlation of some CDRs is also poor, probably due to structures getting included in a cluster that have mis-modeled coordinates, such as peptide flips solved with a bad molecular replacement template. For example, in the L3-9-cis7-1 cluster, we find structures with a cis peptide bond at position 7 that do not have a proline residue at that position, which is almost certainly related to molecular replacement and incorrect modeling. Some clusters also have very similar sequence profiles; in some cases a small cluster is very similar in RMSD space and sequence profile to a very large cluster but has a peptide flip, which alters the ψ and ϕ of two consecutive residues by 180° each (Hayward 2001) without significantly disturbing the positions of neighboring residues. Finally, as we show below, some of the 2011 clusters have poor electron density of the backbone carbonyl atom at individual positions, usually indicating that a flipped peptide has been incorrectly modeled.

With all of these considerations in mind, in this paper we revisit the problem of clustering antibody CDR structures and contemplate the definition of “canonical CDR conformations.” We have utilized several principles to establish more robust CDR clusters than previous efforts:

1. We have established the clusters based on structures that pass an electron density criterion for backbone atoms; for this purpose we use the Electron Density of Individual Atoms (EDIA) (Meyder et al. 2017) over the more traditional B-factor and resolution cutoffs, which do not always correlate with a high-degree of electron density fit to atomic coordinates. We use density-based clustering (DBSCAN) on the high EDIA data (EDIA≥0.7) to define the true clusters from noise, some of which are mismodeled; noise structures occur frequently at low resolution (average EDIA is about 0.8 at 2.8 Å), or because of unusual sequences or engineered mutations, or because of incorrect fitting of electron density, often because of molecular replacement from incorrect templates.
2. We use a maximum dihedral angle metric, which means that the distance between two loop structures is the angular distance function used in directional statistics (D=2(1-cos(θ_1_-θ_2_))) (Mardia and Jupp 2000) for the largest dihedral angle different of ϕ,ψ,ω over all the residues of the loop. This metric clearly separates structures which have peptide flips relative to other structures, which can be missed by the *average* dihedral difference we used previously.
3. We optimized the DBSCAN parameters by identifying consensus clusters produced by DBSCAN over a range of parameters, such that the largest clusters are found without merging unrelated conformations (generally with two peaks of density in the Ramachandran map).
4. To identify likely “canonical” clusters, we established a minimum of 10 unique sequences in the DBSCAN clusters run without an EDIA cutoff (in all X-ray or EM structures with resolution ≤ 3.5 Å), where the clusters are defined in Step 3 but the number of unique sequences comes from corresponding clusters calculated over the whole PDB.
5. We did not cluster CDR lengths (for CDR1, CDR2, CDR4) that do not occur in germline variable gene sequences and which arise only through somatic insertion and deletion (e.g. H1-10, H1-12, L2-6). Unlike the approach of Nowak et al. (Nowak et al. 2016), we did not cluster CDRs in a length-independent manner. Somatic insertions and deletions in CDRs occur but are not common, and the utility of clusters with multiple CDR lengths is not clear.
6. Each cluster at EDIA 0.7 and at 0.0 (no cutoff) had to contain 1% of the chains for that CDR length. This was done to remove some very small clusters that represent very little of the PDB.
7. Some exceptions were made only when these criteria resulted in no clusters for a given CDR length (e.g., H2-11-1, H3-5-2, H4-6-1, H4-7-1, L1-8-1, L3-13-2).

In the new clustering, for the H1, H2, L1, L2, and L3, for which we had 72 clusters in 2011, while we now have 52 clusters, of which 16 are new and 36 are the same as defined in North et al. The total numbers of clusters for each CDR are as follows: H1 (8); H2 (8); H3 (13); H4 (3); L1 (17); L2 (3); L3 (17); L4 (4), for a total of 73 canonical clusters.

## Results

### Issues with clustering of North et al. (2011)

The work is an update of the clustering that we presented in previously (North et al. 2011). Our clustering is widely used to categorize new structures (Teplyakov et al. 2016) and to analyze molecular dynamics simulations of antibodies (Fernández-Quintero et al. 2020). Table 1 shows cluster membership for each of the 72 non-H3 North cluster in the 2011 paper versus September 2022 based on a 40° assignment cutoff to the median of each cluster, i.e. using the same method to assign CDRs to clusters that we have used in our PyIgClassify database (Adolf-Bryfogle et al. 2015). Based on the listings in Table 1, many of the clusters that were initially established during the North et al. work have not grown substantially in the more than 10 years since publication (e.g., H1-13-11, H2-9-2, L1-10-2, L3-11-cis7-1). Additionally, many of the clusters, especially clusters with cis-peptide bonds included, are singleton clusters and have not shown any membership growth. This is a strong motivating factor to revisit the clustering work to ensure that the defined canonical clusters are robust, with each having a significant number of unique sequences and solid experimental support.

**Table 1.** North clusters in 2011 and 2022. CDR clusters which have been deleted either because of low electron density and/or too few sequences are shown in red type.

| Cluster | Chains<br>(2011) | Seqs<br>(2011) | Chains<br>(2022) | Seqs<br>(2022) |
| --- | --- | --- | --- | --- |
| H1-10-1 | 2 | 2 | 0 | 0 |
| H1-12-1 | 1 | 1 | 0 | 0 |
| H1-13-1 | 267 | 213 | 8117 | 1642 |
| H1-13-2 | 7 | 7 | 200 | 94 |
| H1-13-3 | 5 | 5 | 248 | 99 |
| H1-13-4 | 4 | 4 | 289 | 75 |
| H1-13-5 | 4 | 4 | 227 | 63 |
| H1-13-6 | 4 | 4 | 40 | 18 |
| H1-13-7 | 3 | 3 | 116 | 35 |
| H1-13-8 | 3 | 3 | 58 | 6 |
| H1-13-9 | 3 | 3 | 16 | 7 |
| H1-13-10 | 2 | 2 | 23 | 10 |
| H1-13-11 | 2 | 2 | 8 | 3 |
| H1-13-cis9-1 | 2 | 1 | 8 | 1 |
| H1-14-1 | 11 | 7 | 193 | 48 |
| H1-15-1 | 9 | 7 | 324 | 93 |
| H1-16-1 | 1 | 1 | 0 | 0 |
| H2-8-1 | 2 | 2 | 0 | 0 |
| H2-9-1 | 77 | 57 | 2187 | 550 |
| H2-9-2 | 2 | 2 | 16 | 6 |
| H2-9-3 | 2 | 2 | 129 | 25 |
| H2-10-1 | 155 | 131 | 4150 | 1020 |
| H2-10-2 | 42 | 40 | 2528 | 612 |
| H2-10-3 | 11 | 9 | 234 | 77 |
| H2-10-4 | 7 | 7 | 280 | 34 |
| H2-10-5 | 3 | 2 | 111 | 38 |
| H2-10-6 | 3 | 3 | 304 | 93 |
| H2-10-7 | 2 | 2 | 12 | 8 |
| H2-10-8 | 2 | 2 | 25 | 13 |
| H2-10-9 | 2 | 2 | 37 | 19 |
| H2-12-1 | 26 | 22 | 488 | 95 |
| H2-15-1 | 1 | 1 | 0 | 0 |
| L1-10-1 | 20 | 16 | 263 | 61 |
| L1-10-2 | 2 | 2 | 104 | 4 |
| L1-11-1 | 76 | 57 | 2878 | 440 |
| L1-11-2 | 55 | 37 | 897 | 194 |
| L1-11-3 | 5 | 5 | 281 | 85 |
| L1-12-1 | 5 | 5 | 449 | 89 |
| L1-12-2 | 5 | 5 | 261 | 40 |
| L1-12-3 | 2 | 2 | 61 | 11 |
| L1-13-1 | 7 | 7 | 505 | 97 |
| L1-13-2 | 4 | 4 | 137 | 16 |
| L1-14-1 | 14 | 8 | 264 | 27 |
| L1-14-2 | 4 | 4 | 350 | 74 |
| L1-15-1 | 11 | 9 | 452 | 99 |
| L1-15-2 | 2 | 2 | 13 | 7 |
| L1-16-1 | 68 | 50 | 1029 | 192 |
| L1-17-1 | 21 | 17 | 549 | 111 |
| L2-8-1 | 290 | 159 | 8371 | 988 |
| L2-8-2 | 9 | 8 | 482 | 148 |
| L2-8-3 | 3 | 2 | 53 | 13 |
| L2-8-4 | 2 | 2 | 401 | 99 |
| L2-8-5 | 2 | 2 | 295 | 76 |
| L2-12-1 | 2 | 1 | 13 | 6 |
| L2-12-2 | 2 | 1 | 57 | 11 |
| L3-7-1 | 2 | 2 | 0 | 0 |
| L3-8-1 | 15 | 13 | 335 | 74 |
| L3-8-2 | 4 | 4 | 25 | 16 |
| L3-8-cis6-1 | 3 | 2 | 5 | 3 |
| L3-9-1 | 22 | 17 | 347 | 69 |
| L3-9-2 | 12 | 12 | 587 | 167 |
| L3-9-cis6-1 | 1 | 1 | 5 | 2 |
| L3-9-cis7-1 | 219 | 182 | 4678 | 907 |
| L3-9-cis7-2 | 8 | 7 | 65 | 25 |
| L3-10-1 | 6 | 5 | 216 | 53 |
| L3-10-cis7and8-1 | 1 | 1 | 54 | 23 |
| L3-10-cis8-1 | 2 | 2 | 2 | 1 |
| L3-11-1 | 9 | 9 | 501 | 136 |
| L3-11-cis7-1 | 1 | 1 | 61 | 3 |
| L3-12-1 | 1 | 1 | 0 | 0 |
| L3-13-1 | 3 | 2 | 21 | 6 |

In order to build high-confidence datasets to determine conformational families, we used the EDIA software from Meyder et al. (Meyder et al. 2017). EDIA calculates the fit of atomic coordinates to the local electron density within a sphere surrounding the atom in question. EDIA is a rigorous method for determining the fit of atomic coordinates and their local density, and determines a cutoff for the EDIA score of 0.8 for which researchers should take particular caution in considering the structures that those atoms represent. EDIA is highly dependent on resolution with high-resolution structures (<1.5 Å) having average EDIA scores of about 1.0 for backbone atoms, while low resolution structures around 2.6 Å have an average minimum backbone EDIA score per residue of around 0.8 (Figure 1). Since the distribution of resolution across the members of different CDR clusters may differ, thus affecting the average at each position, it is *variation* of the average across the residues in one cluster that may be indicative of poor fitting of electron density at one or more residues. In Figure 2, in the left column we show several North CDR clusters which have generally uniform mean values of EDIA across the length of each CDR. In the right column in Figure 2, we show examples of North clusters with fluctuating EDIA mean values for the same CDR lengths shown in the left column. While EDIA is strongly dependent on resolution, some clusters have individual residues that have lower average EDIA scores than the rest of the CDR, indicating potential misfitting of these coordinates. For example, H1-13-2 has lower EDIA values at positions 3 and 4 compared to the remaining residues in the CDR, while the residues in the much larger cluster, H1-13-1, has consistent EDIA distributions across the length of the CDR. The structure of H1-13-2 is very similar to that of H1-13-1, except at positions 3 and 4 (marked with red arrows in Figure 1), where H1-13-2 (Ramachandran string `BBAABBAAABBBB`) has a “peptide flip” from the structure of H1-13-1 (Ramachandran string `BBBLBBAAABBBB`). In a peptide flip between two structures, the ψ of residue *N* and ϕ of residue *N*+1 of one structure both differ by ~180° from the same values in the other structure (Hayward 2001), displacing the oxygen atom of residue *N* by about 3 Å. So for H1-13-1 to H1-13-2 there is a flip of “BL” to “AA,” which is the most common peptide flip transition in loop structures (first residue, B→A is a 180° change in ψ; second residue, L→A is a 180° change in ϕ). Peptide flips also occur between clusters H2-10-4 (`BBBBLLABBB`, correctly modeled) and H2-10-7 (`BBBBEAABBB`, incorrectly modeled), and between L3-8-1 (`BBAABEBB`) and L3-8-2 (`BBABEBBB`). Here again these are peptide flips: LL→EA is a 180° change in ψ in the first position (L→E) and a 180° change in ϕ at second position (L→A); AB→BE is a 180° change in ψ in the first position (A→B) and a 180° change in ϕ at second position (B→E). A comparison of H1-13-1 and H1-13-2 structures showing the poor electron density at these residues in H1-13-2 is shown in Figure 3.

**Figure 1.**
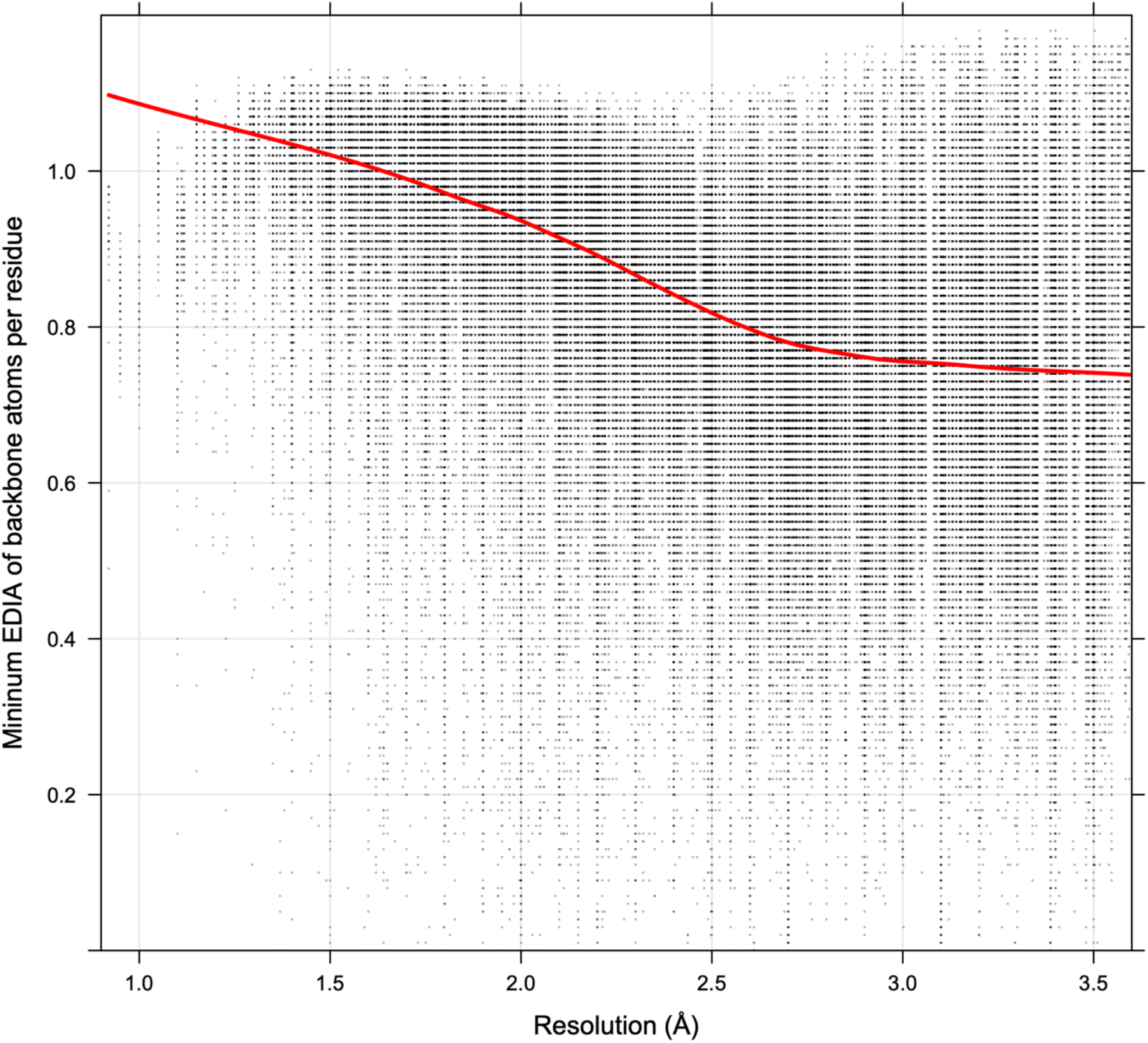
Minimum EDIA values over backbone atoms for individual residues in the full CDR data set vs resolution. The red line represents a Loess regression of the EDIA values.

**Figure 2.**
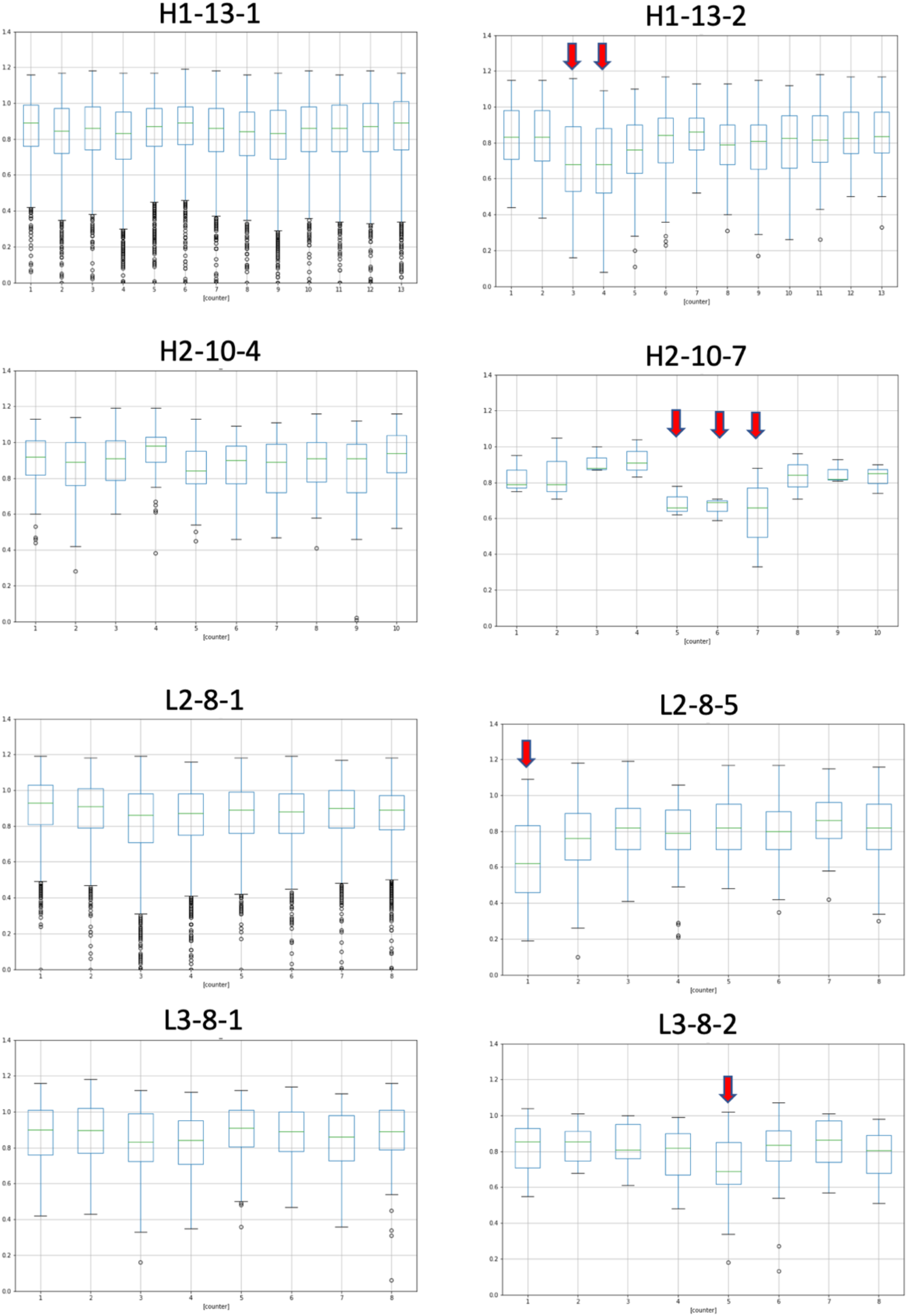
EDIA distributions for stable clusters (left column) and unstable clusters (right column). Red arrows in the right column indicate low electron density positions usually associated with incorrectly modeled peptide flips relative to the cluster in the left column. Clusters in the right column contain peptide flips relative to the much larger clusters in the left column. First row: H1-13-1 (Ramachandran string BB**BL**BBAAABBBB) → H1-13-2 (BB**AA**BBAAABBBB. Second row: H2-10-4 (BBBB**LL**ABBB) → H2-10-7 (BBBB**EA**ABBB). Third row: L2-8-5 (**AB**AABBBB) is a peptide flip from L2-8-4 (**BE**AABBBB) (not shown). Fourth row: L3-8-1 (BBA**AB**EBB) → L3-8-2 (BBA**BE**BBB).

**Figure 3.**
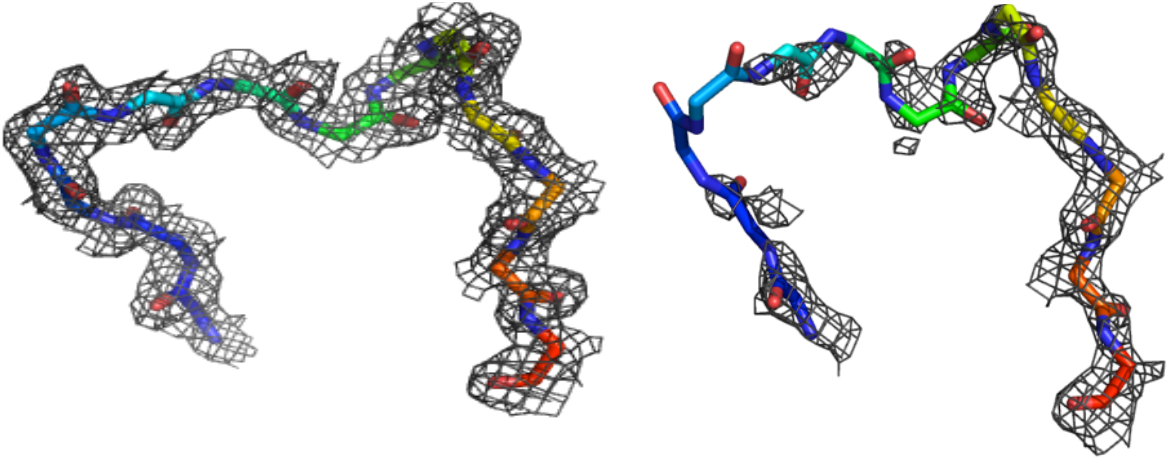
Electron density for representatives of North clusters H1-13-1 (left) and H1-13-2 (right), showing low electron density at positions 3 and 4 in H1-13-2 (an BL→AA peptide flip, incorrectly modeled in most North cluster H1-13-2 structures).

### Clusters from all EDIA cutoff datasets

In this work, we incorporate the EDIA score into our structure quality assessment by generating multiple datasets, where atoms for a chosen all backbone atoms of a CDR must meet a particular minimum EDIA score cutoff in order to be considered in the clustering set for that particular CDR-length. Specifically, we generated data sets of X-ray structures with EDIA cutoffs of 0.1, 0.2, 0.3, 0.4, 0.5, 0.6, 0.7, 0.8, and 0.9, and a tenth data set consisting of all CDR X-ray and EM structures in the PDB with resolution ≤ 3.5 Å (“EDIA=0.0”, i.e., no cutoff). For the EDIA cutoff datasets, only X-ray structures that had deposited structure factor files in the PDB were included in the analysis. For the PDB-wide analysis (EDIA=0.0), we included structures that did not have structure factor files. Table 2 shows the number of structures at each EDIA cutoff. Since EDIA is dependent on resolution and the average backbone atom in a 2.6 Å structure has an EDIA of 0.8 (Figure 1), there is a steep decline in the amount of data at EDIA values above 0.6.

**Table 2.** Data sets for clustering at different minimum EDIA cutoff values.

| CDR | EDIA<br>Cutoff | Number of<br>ordered CDRs | CDR | EDIA<br>Cutoff | Number of<br>ordered CDRs |
| --- | --- | --- | --- | --- | --- |
| H1 | 0.0 | 8948 | L1 | 0.0 | 7566 |
| H1 | 0.1 | 5026 | L1 | 0.1 | 4504 |
| H1 | 0.2 | 4945 | L1 | 0.2 | 4445 |
| H1 | 0.3 | 4781 | L1 | 0.3 | 4293 |
| H1 | 0.4 | 4488 | L1 | 0.4 | 4010 |
| H1 | 0.5 | 3843 | L1 | 0.5 | 3425 |
| H1 | 0.6 | 2778 | L1 | 0.6 | 2520 |
| H1 | 0.7 | 1743 | L1 | 0.7 | 1598 |
| H1 | 0.8 | 1086 | L1 | 0.8 | 956 |
| H1 | 0.9 | 482 | L1 | 0.9 | 451 |
| H2 | 0.0 | 9065 | L2 | 0.0 | 7631 |
| H2 | 0.1 | 5133 | L2 | 0.1 | 4563 |
| H2 | 0.2 | 5085 | L2 | 0.2 | 4552 |
| H2 | 0.3 | 4978 | L2 | 0.3 | 4486 |
| H2 | 0.4 | 4726 | L2 | 0.4 | 4326 |
| H2 | 0.5 | 4201 | L2 | 0.5 | 3943 |
| H2 | 0.6 | 3228 | L2 | 0.6 | 3245 |
| H2 | 0.7 | 2149 | L2 | 0.7 | 2285 |
| H2 | 0.8 | 1313 | L2 | 0.8 | 1441 |
| H2 | 0.9 | 666 | L2 | 0.9 | 784 |
| H3 | 0.0 | 8771 | L3 | 0.0 | 7634 |
| H3 | 0.1 | 4894 | L3 | 0.1 | 4549 |
| H3 | 0.2 | 4782 | L3 | 0.2 | 4506 |
| H3 | 0.3 | 4609 | L3 | 0.3 | 4403 |
| H3 | 0.4 | 4275 | L3 | 0.4 | 4156 |
| H3 | 0.5 | 3581 | L3 | 0.5 | 3645 |
| H3 | 0.6 | 2614 | L3 | 0.6 | 2812 |
| H3 | 0.7 | 1728 | L3 | 0.7 | 1878 |
| H3 | 0.8 | 1078 | L3 | 0.8 | 1200 |
| H3 | 0.9 | 547 | L3 | 0.9 | 664 |
| H4 | 0.0 | 9015 | L4 | 0.0 | 7601 |
| H4 | 0.1 | 5101 | L4 | 0.1 | 4540 |
| H4 | 0.2 | 5056 | L4 | 0.2 | 4515 |
| H4 | 0.3 | 4966 | L4 | 0.3 | 4451 |
| H4 | 0.4 | 4792 | L4 | 0.4 | 4304 |
| H4 | 0.5 | 4352 | L4 | 0.5 | 3950 |
| H4 | 0.6 | 3494 | L4 | 0.6 | 3250 |
| H4 | 0.7 | 2379 | L4 | 0.7 | 2240 |
| H4 | 0.8 | 1386 | L4 | 0.8 | 1261 |
| H4 | 0.9 | 606 | L4 | 0.9 | 528 |

We work through an example of CDR-length L3-8 to help illustrate the data. The clustering with DBSCAN for L3-8 at different EDIA cutoffs is shown in Figure 4. As the EDIA cutoff increases, the number of clusters decreases. Clusters that arise in the calculations on different EDIA data sets are compared using a Simpson index metric (see Methods) that determines whether there is significant overlap. Typically, the clusters at higher EDIA cutoffs are subsets of clusters from the larger data sets created at lower EDIA cutoffs. An exact subset has a Simpson index of 1.0. The original North cluster L3-8-2 disappears at EDIA above 0.5 and above. The new clusters, L3-8-3 and L3-8-4, are stable until EDIA values of 0.8 and 0.7 respectively. The EDIA distributions for the 0.0 cutoff are shown in Figure 5. While the differences are not dramatic, L3-8-2 has lower EDIA values and a larger variance at position 4. The Ramachandran plots show that L3-8-2 is a peptide flip (BA → AL) of the (new) cluster L3-8-4 at positions 4-5.

**Figure 4.**
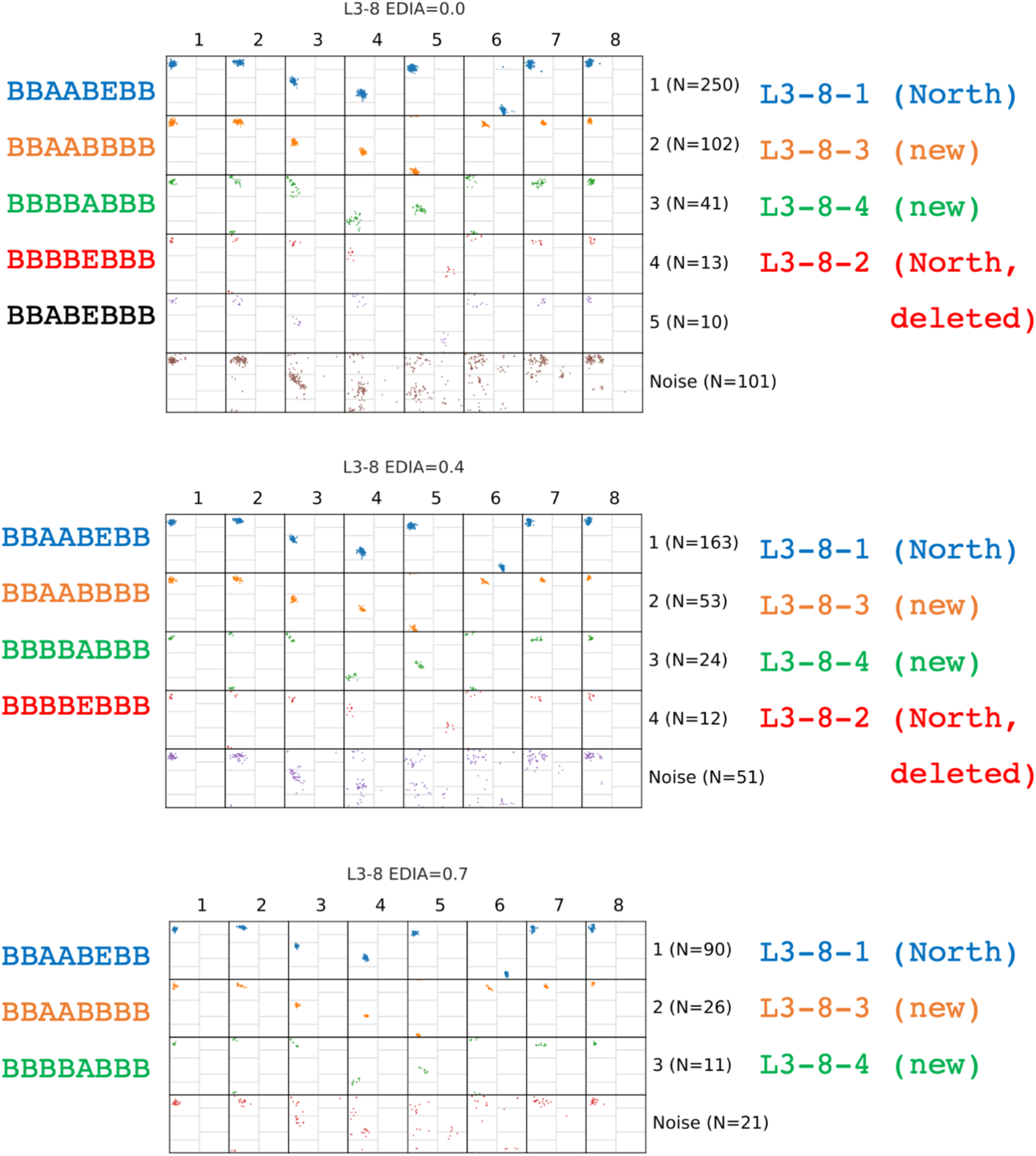
Ramachandran plots for DBSCAN clustering for CDR L3-8 at EDIA cutoff values of 0.0, 0.4, and 0.7. The Ramachandran strings for each cluster are shown at left (A=alpha region; B=beta region; L=left-handed region; E=epsilon region (lower right and far upper right region of Ramachandran maps. The borders of each region are shown in thin gray lines in each plot. The cluster names are shown at right. L3-8-1 is a preserved North cluster and L3-8-2 is deleted from the new clustering. The fifth cluster at EDIA=0.0 is not maintained at higher EDIA cutoffs and is skipped in the new clustering. To maintain compatibility with the North clustering, the name L3-8-2 is retired and the new clusters are given previously unused names, L3-8-3 (orange) and L3-8-4 (green).

**Figure 5.**
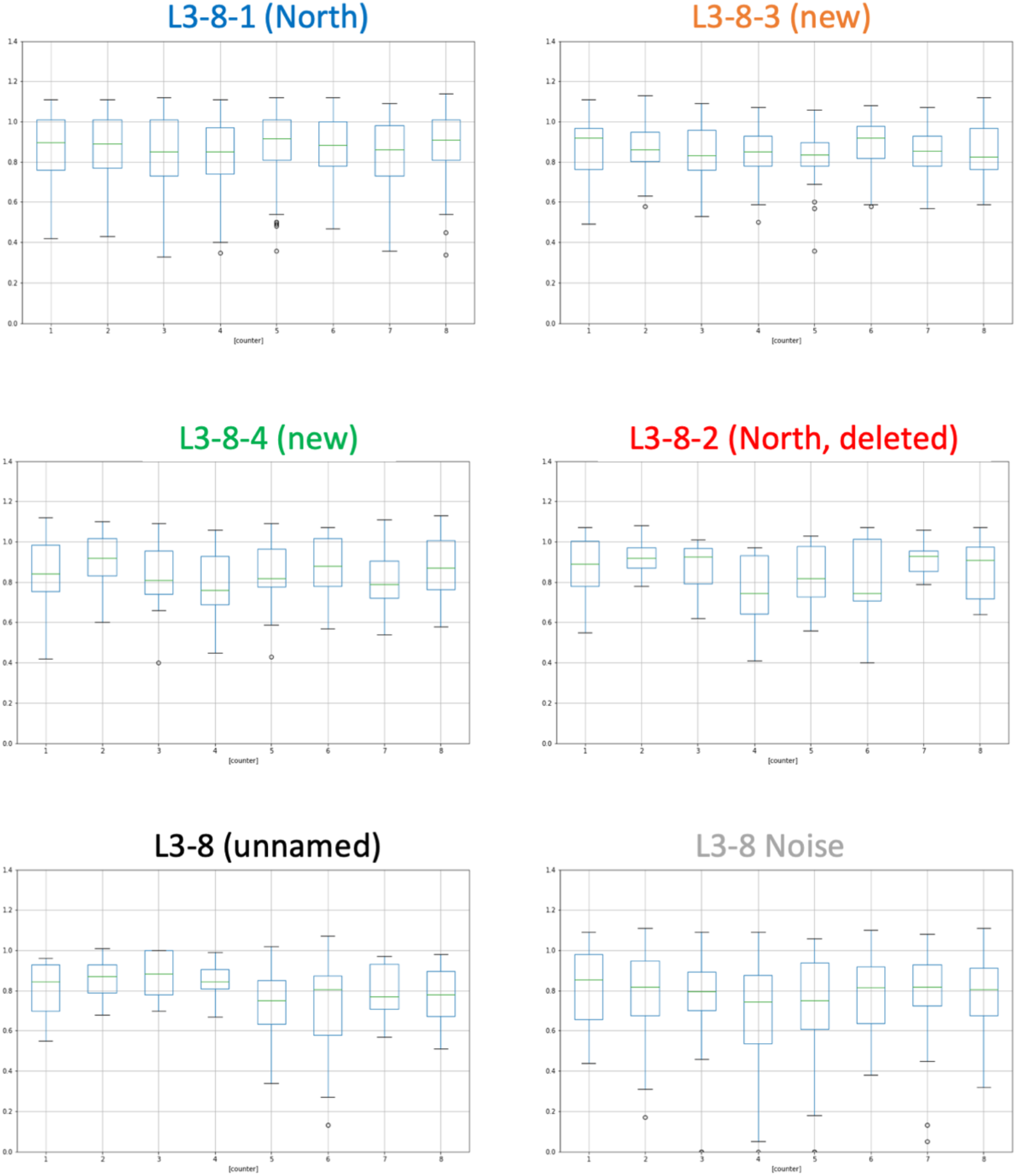
EDIA distributions for all 5 clusters determined from DBSCAN applied to L3-8 EDIA=0.0 data (no cutoff) plus the noise data. The L3-8-2 (North) cluster is not observed in data with EDIA cutoff higher than 0.5.

After examination of the data for the clusters of all CDR lengths, we chose rules designed to identify “canonical clusters” – those with enough structures with sufficient electron densities and enough sequences across the PDB. The cutoffs are somewhat arbitrary but seem reasonable in view of the data. The criteria are as follows:

1. There must be a cluster at EDIA cutoff as high as 0.7 or higher
2. The clusters at EDIA of 0.0 and at 0.7 must contain at least 1% of the chains clustered in those data sets to eliminate many small clusters for common CDR lengths (such as H1-13).
3. There must be at least 10 unique sequences in the EDIA=0.0 cluster.
4. Exceptions to these rules were made in some cases if no clusters resulted but a minimum of 5 unique sequences was required in the EDIA=0.0 cluster (e.g. H2-11-1).

Examples of clusters that we kept and examples of those that were deleted (if they were present in the North clustering) or skipped (if they were not present in the North clustering) are shown in Tables 3 and 4 for the light and heavy chains. Each of the lengths shown (L1-11, L2-8, L3-9, L4-6, H1-13, H2-10, H3-12, H4-8) are the most common lengths for each of the 8 CDRs. The extensive data in these tables show that for most of the commonest clusters, the percentage of chains in the cluster rises as a function of the EDIA cutoff. For example, for H1-13-1, with no EDIA cutoff, the cluster represents 64% of the H1-13 CDRs but at a cutoff of 0.9, H1-13-1 represents 86% of the chains in the data set. The percentage of chains that end up placed in noise by our grid-DBSCAN algorithm decreases as a function of increasing EDIA in most cases. For example, for L2-8, the percentage of chains in noise for the EDIA=0.0 data set is 20.3% while for the EDIA=0.9 data set, noise represents 4.4% of the data. The clusters listed that were deleted (if present in the North clustering) or skipped (if not present) generally show flat or decreasing representation as EDIA increases. The reason for not including them in the final list of clusters is given in the second column, indicating whether it is either missing clusters at higher EDIA, insufficient chains at EDIA=0.0 or EDIA=0.7, and/or insufficient unique sequences at EDIA=0.0.

**Table 3.** Number of sequences, domains, and percentage of CDR lengths for selected light-chain clusters.

| Cluster | Status | EDIA | Ntotal<br>Chains<br>(CDRlength) | Nseq | Ndomains | %Chains<br>(CDRlength) | Nseq<br>(Noise) | Nchains<br>(Noise) | %Chains<br>(Noise) |
| --- | --- | --- | --- | --- | --- | --- | --- | --- | --- |
| L1-11-1 | Keep | 0.0 | 3402 | 351 | 1838 | 54.0 | 134 | 384 | 11.3 |
|  |  | 0.1 | 1818 | 229 | 996 | 54.8 | 79 | 194 | 10.7 |
|  |  | 0.2 | 1808 | 229 | 999 | 55.3 | 78 | 181 | 10.0 |
|  |  | 0.3 | 1767 | 226 | 980 | 55.5 | 69 | 169 | 9.6 |
|  |  | 0.4 | 1665 | 225 | 953 | 57.2 | 60 | 146 | 8.8 |
|  |  | 0.5 | 1478 | 215 | 863 | 58.4 | 69 | 144 | 9.7 |
|  |  | 0.6 | 1113 | 191 | 681 | 61.2 | 48 | 75 | 6.7 |
|  |  | 0.7 | 738 | 153 | 471 | 63.8 | 30 | 49 | 6.6 |
|  |  | 0.8 | 437 | 115 | 280 | 64.1 | 17 | 22 | 5.0 |
|  |  | 0.9 | 216 | 75 | 141 | 65.3 | 21 | 22 | 10.2 |
| L1-11-a | Skip<br>(maxEDIA=0.3;<br>Nseq<10) | 0.0 | 3402 | 3 | 26 | 0.8 | 134 | 384 | 11.3 |
|  |  | 0.1 | 1818 | 3 | 14 | 0.8 | 79 | 194 | 10.7 |
|  |  | 0.2 | 1808 | 3 | 14 | 0.8 | 78 | 181 | 10.0 |
|  |  | 0.3 | 1767 | 3 | 14 | 0.8 | 69 | 169 | 9.6 |
| L2-8-1 | Keep | 0.0 | 7555 | 782 | 5129 | 67.9 | 382 | 1533 | 20.3 |
|  |  | 0.1 | 4500 | 591 | 3446 | 76.6 | 234 | 706 | 15.7 |
|  |  | 0.2 | 4490 | 591 | 3442 | 76.7 | 232 | 703 | 15.7 |
|  |  | 0.3 | 4430 | 590 | 3426 | 77.3 | 230 | 676 | 15.3 |
|  |  | 0.4 | 4274 | 588 | 3367 | 78.8 | 212 | 620 | 14.5 |
|  |  | 0.5 | 3896 | 579 | 3148 | 80.8 | 195 | 513 | 13.2 |
|  |  | 0.6 | 3208 | 560 | 2831 | 88.3 | 99 | 180 | 5.6 |
|  |  | 0.7 | 2266 | 484 | 2085 | 92.0 | 43 | 74 | 3.3 |
|  |  | 0.8 | 1429 | 361 | 1310 | 91.7 | 54 | 75 | 5.3 |
|  |  | 0.9 | 777 | 238 | 723 | 93.1 | 28 | 34 | 4.4 |
| L2-8-5 | Delete<br>(maxEDIA=0.4) | 0.0 | 7555 | 29 | 90 | 1.2 | 382 | 1533 | 20.3 |
|  |  | 0.1 | 4500 | 13 | 37 | 0.8 | 234 | 706 | 15.7 |
|  |  | 0.2 | 4490 | 14 | 37 | 0.8 | 232 | 703 | 15.7 |
|  |  | 0.3 | 4430 | 12 | 32 | 0.7 | 230 | 676 | 15.3 |
|  |  | 0.4 | 4274 | 11 | 23 | 0.5 | 212 | 620 | 14.5 |
| L3-9-cis7-1 | Keep | 0.0 | 5033 | 811 | 3553 | 70.6 | 245 | 525 | 10.4 |
|  |  | 0.1 | 2766 | 547 | 2003 | 72.4 | 133 | 232 | 8.4 |
|  |  | 0.2 | 2750 | 546 | 2000 | 72.7 | 132 | 228 | 8.3 |
|  |  | 0.3 | 2695 | 536 | 1940 | 72.0 | 148 | 252 | 9.4 |
|  |  | 0.4 | 2569 | 539 | 1899 | 73.9 | 113 | 193 | 7.5 |
|  |  | 0.5 | 2292 | 521 | 1719 | 75.0 | 106 | 173 | 7.6 |
|  |  | 0.6 | 1758 | 454 | 1342 | 76.3 | 94 | 133 | 7.6 |
|  |  | 0.7 | 1157 | 364 | 910 | 78.7 | 60 | 74 | 6.4 |
|  |  | 0.8 | 745 | 283 | 607 | 81.5 | 44 | 59 | 7.9 |
|  |  | 0.9 | 426 | 187 | 335 | 78.6 | 51 | 61 | 14.3 |
| L3-9-cis7-3 | Delete<br>(%chains<1.0) | 0.0 | 5033 | 20 | 47 | 0.9 | 245 | 525 | 10.4 |
|  |  | 0.1 | 2766 | 11 | 24 | 0.9 | 133 | 232 | 8.4 |
|  |  | 0.2 | 2750 | 11 | 24 | 0.9 | 132 | 228 | 8.3 |
|  |  | 0.3 | 2695 | 11 | 24 | 0.9 | 148 | 252 | 9.4 |
|  |  | 0.4 | 2569 | 11 | 23 | 0.9 | 113 | 193 | 7.5 |
|  |  | 0.5 | 2292 | 8 | 17 | 0.7 | 106 | 173 | 7.6 |
|  |  | 0.6 | 1758 | 7 | 17 | 1.0 | 94 | 133 | 7.6 |
|  |  | 0.7 | 1157 | 6 | 13 | 1.1 | 60 | 74 | 6.4 |
| L4-6-1 | Keep | 0.0 | 7464 | 80 | 4556 | 61.0 | 99 | 943 | 12.6 |
|  |  | 0.1 | 4420 | 62 | 2504 | 56.7 | 68 | 501 | 11.3 |
|  |  | 0.2 | 4397 | 62 | 2495 | 56.7 | 69 | 515 | 11.7 |
|  |  | 0.3 | 4335 | 62 | 2474 | 57.1 | 67 | 461 | 10.6 |
|  |  | 0.4 | 4190 | 61 | 2421 | 57.8 | 65 | 419 | 10.0 |
|  |  | 0.5 | 3839 | 59 | 2245 | 58.5 | 61 | 356 | 9.3 |
|  |  | 0.6 | 3154 | 55 | 1881 | 59.6 | 57 | 249 | 7.9 |
|  |  | 0.7 | 2169 | 49 | 1335 | 61.6 | 43 | 98 | 4.5 |
|  |  | 0.8 | 1220 | 42 | 752 | 61.6 | 21 | 40 | 3.3 |
|  |  | 0.9 | 511 | 29 | 308 | 60.3 | 14 | 23 | 4.5 |
| L4-6-a | Skip<br>(maxEDIA=0.5;<br>Nseq<10) | 0.0 | 7464 | 5 | 50 | 0.7 | 99 | 943 | 12.6 |
|  |  | 0.1 | 4420 | 3 | 11 | 0.3 | 68 | 501 | 11.3 |
|  |  | 0.2 | 4397 | 3 | 11 | 0.3 | 69 | 515 | 11.7 |
|  |  | 0.3 | 4335 | 3 | 11 | 0.3 | 67 | 461 | 10.6 |
|  |  | 0.4 | 4190 | 3 | 10 | 0.2 | 65 | 419 | 10.0 |
|  |  | 0.5 | 3839 | 3 | 10 | 0.3 | 61 | 356 | 9.3 |

**Table 4.**
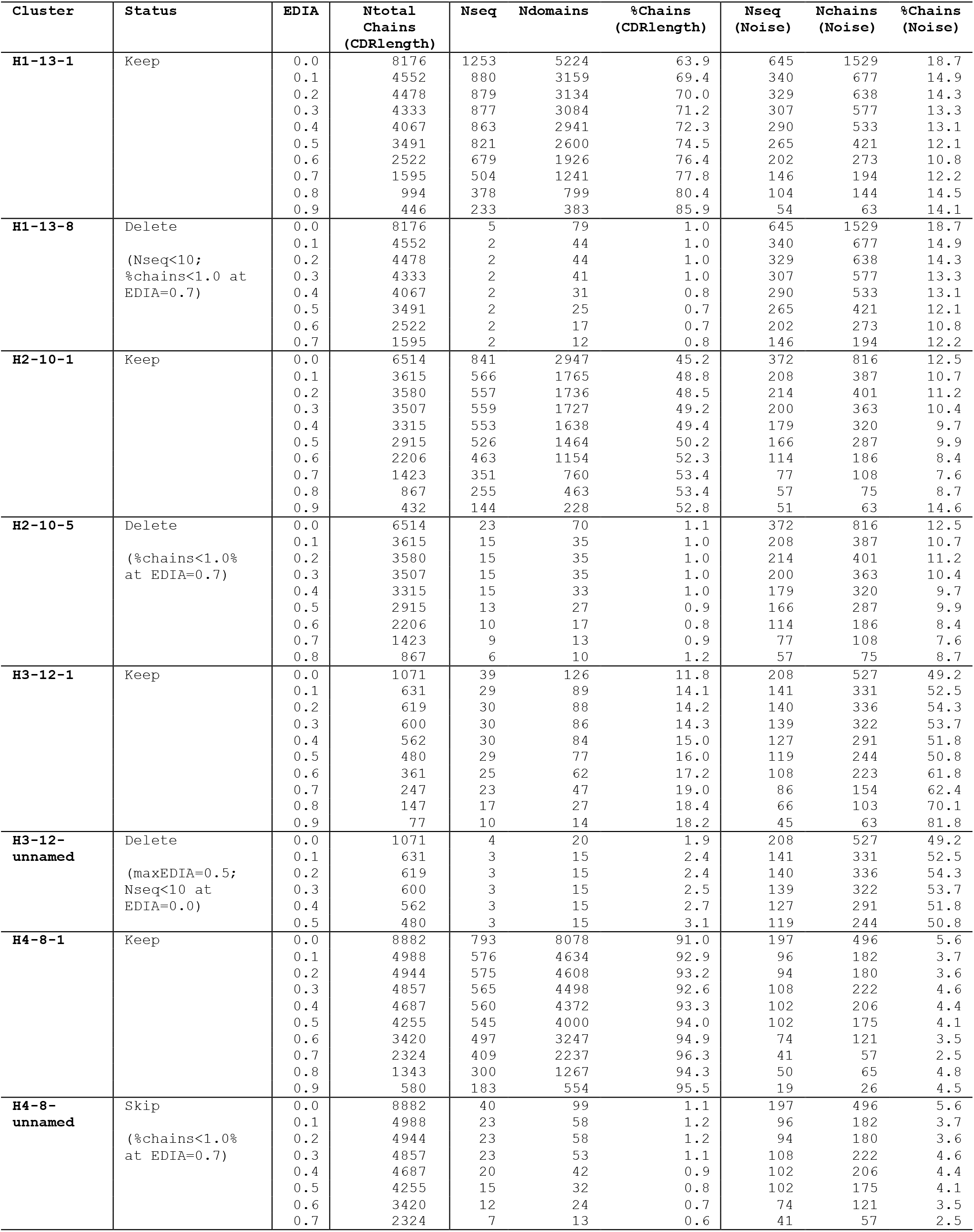
Number of sequences, domains, and percentage of CDR lengths for selected heavy-chain clusters.

### Final clusters

After applying the rules described above, Ramachandran maps for the clusters resulting from DBSCAN at a cutoff of EDIA=0.0 are shown in Figure 6 and logos for their sequence profiles and Ramachandran regions are shown in Figure 7. The EDIA distributions for each extant cluster follow in Figure 8. As shown in Figure 7, as expected, the L region of the Ramachandran map is dominated by Gly, Asn, Asp, and Ser, while the E region is mostly restricted to Gly. Pro is restricted to the A and B regions, but also restricts the Ramachandran region of the preceding residue to the B region, since clashes occur in both the L and A regions with the Cδ atom of trans Pro or the Cα atom in cis-Pro (Ting et al. 2010). Other correlations are evident and can be analyzed further.

**Figure 6.**
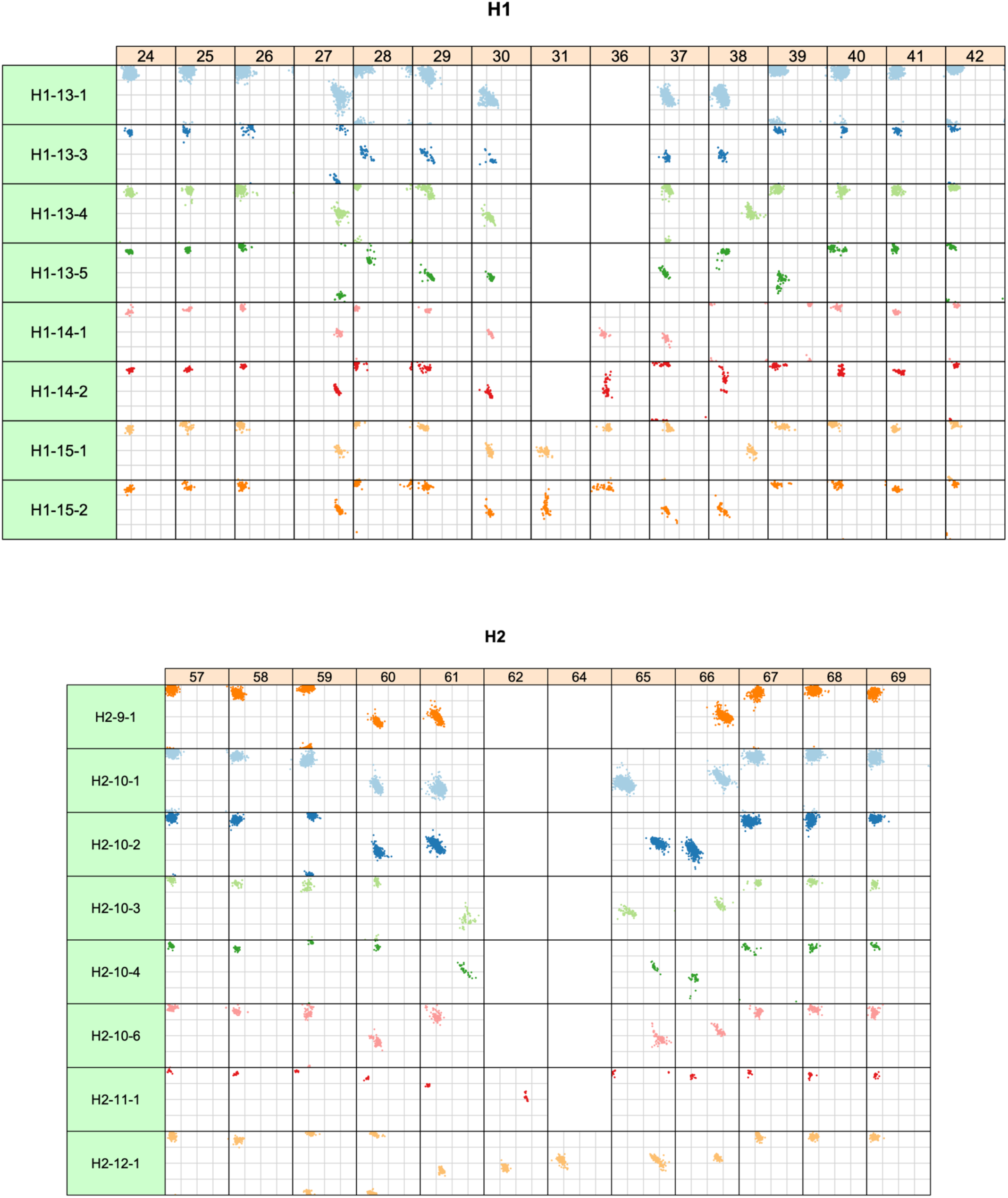
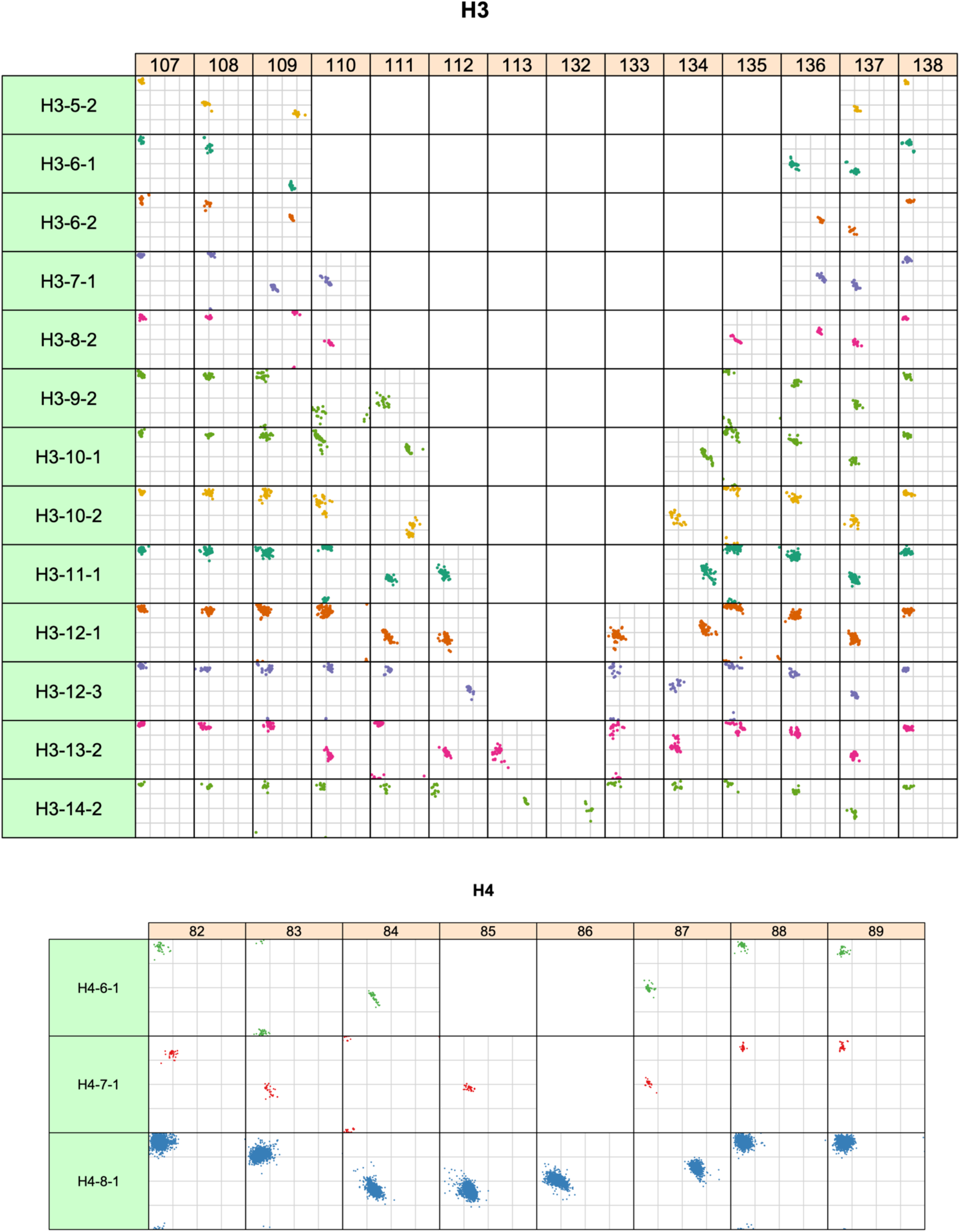
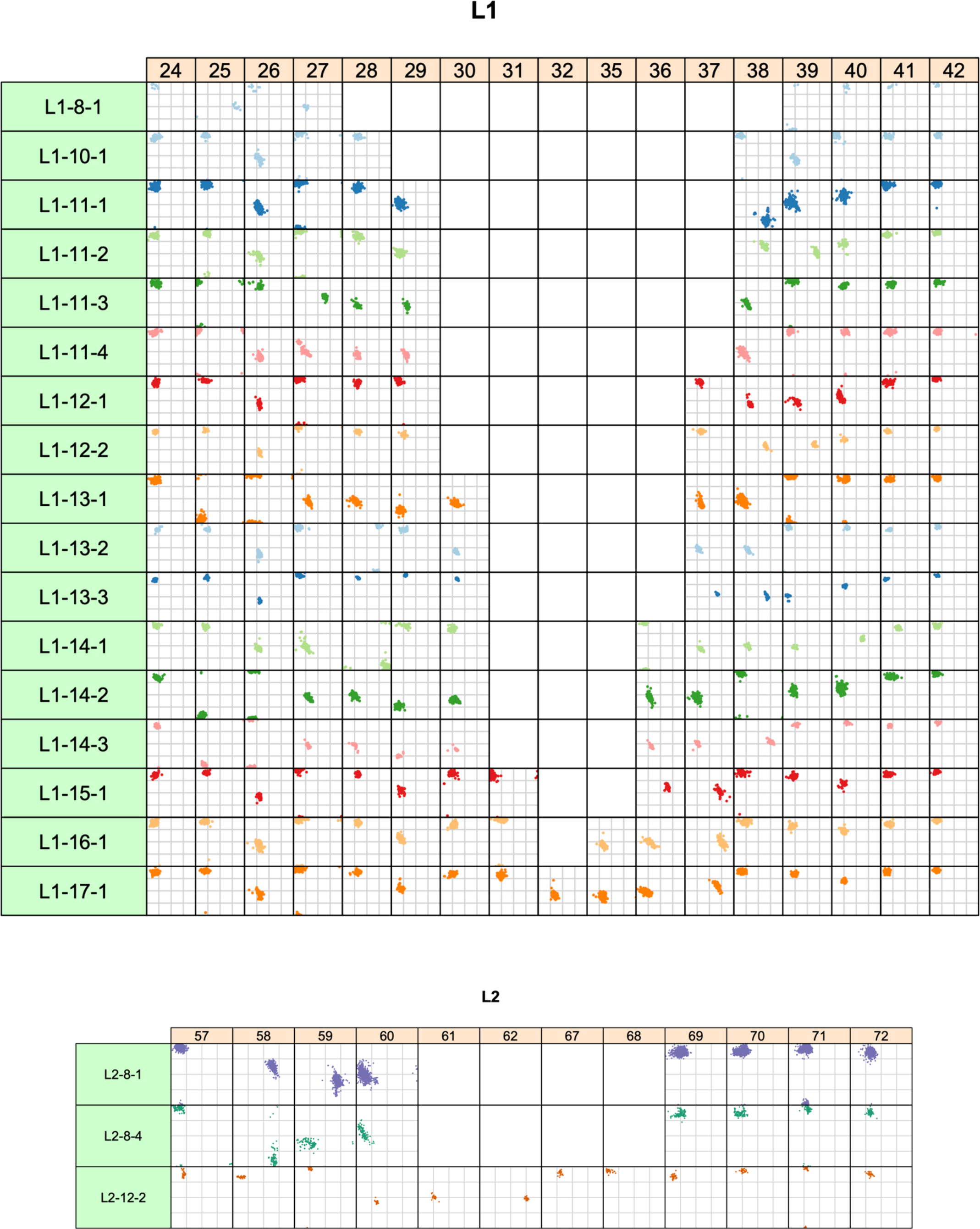
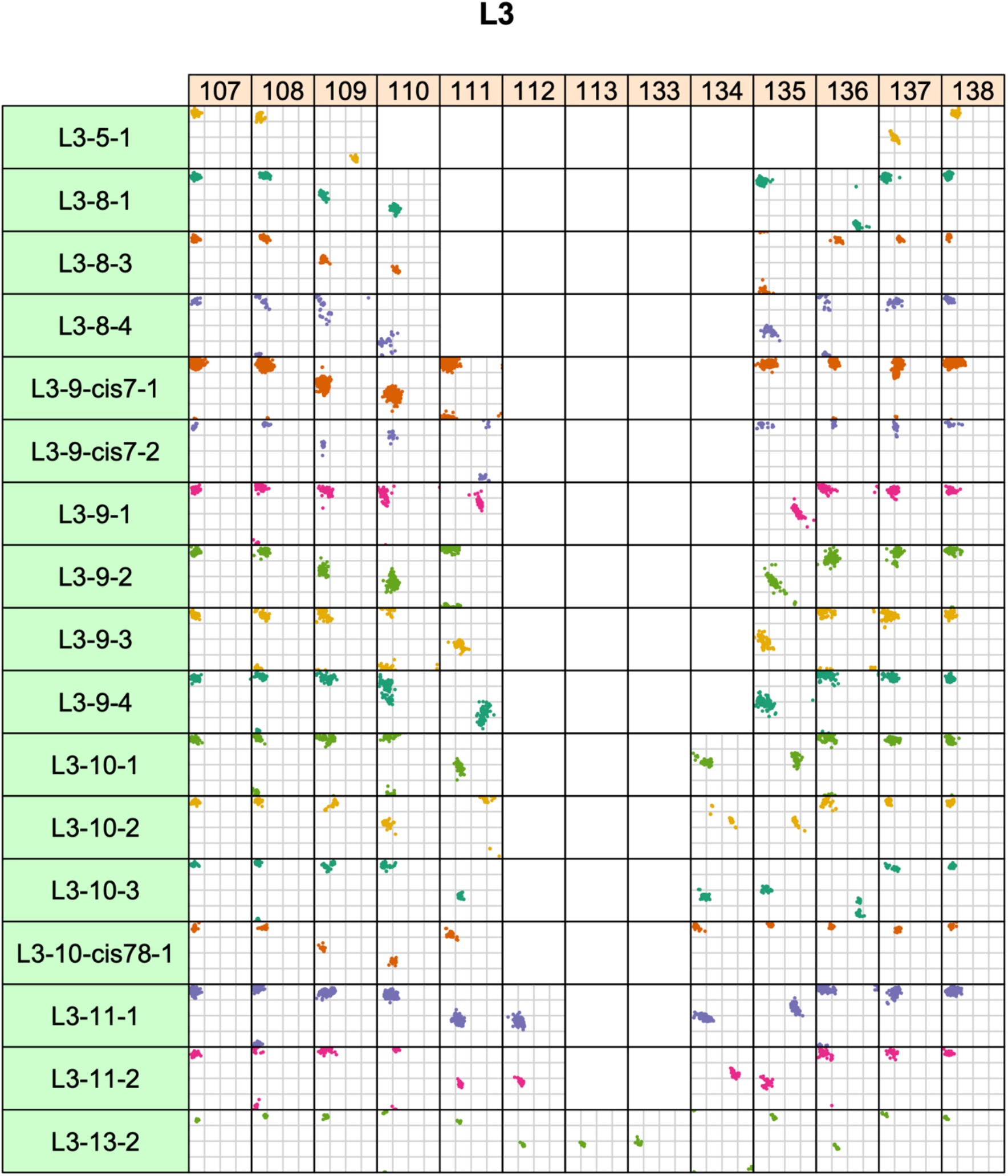
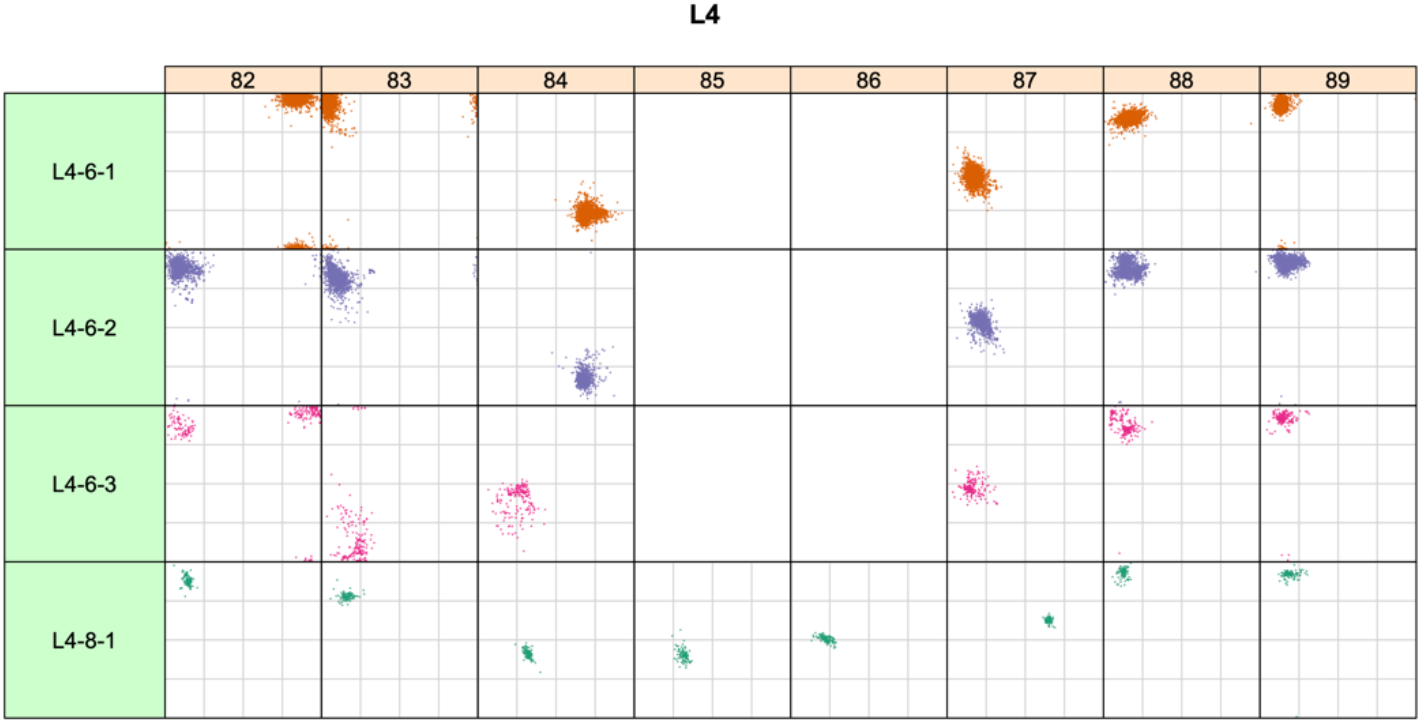
Ramachandran plots for all CDR clusters in the 2022 clustering. The light-gray grid lines are at ϕ and ψ at −90°, 0°, and +90°. Cluster names are backwards compatible with the North et al. clustering. Any previous cluster name that is no longer has been “retired.” Any new clusters have been given names that have not been previously used in the North et al. clustering.

**Figure 7.**
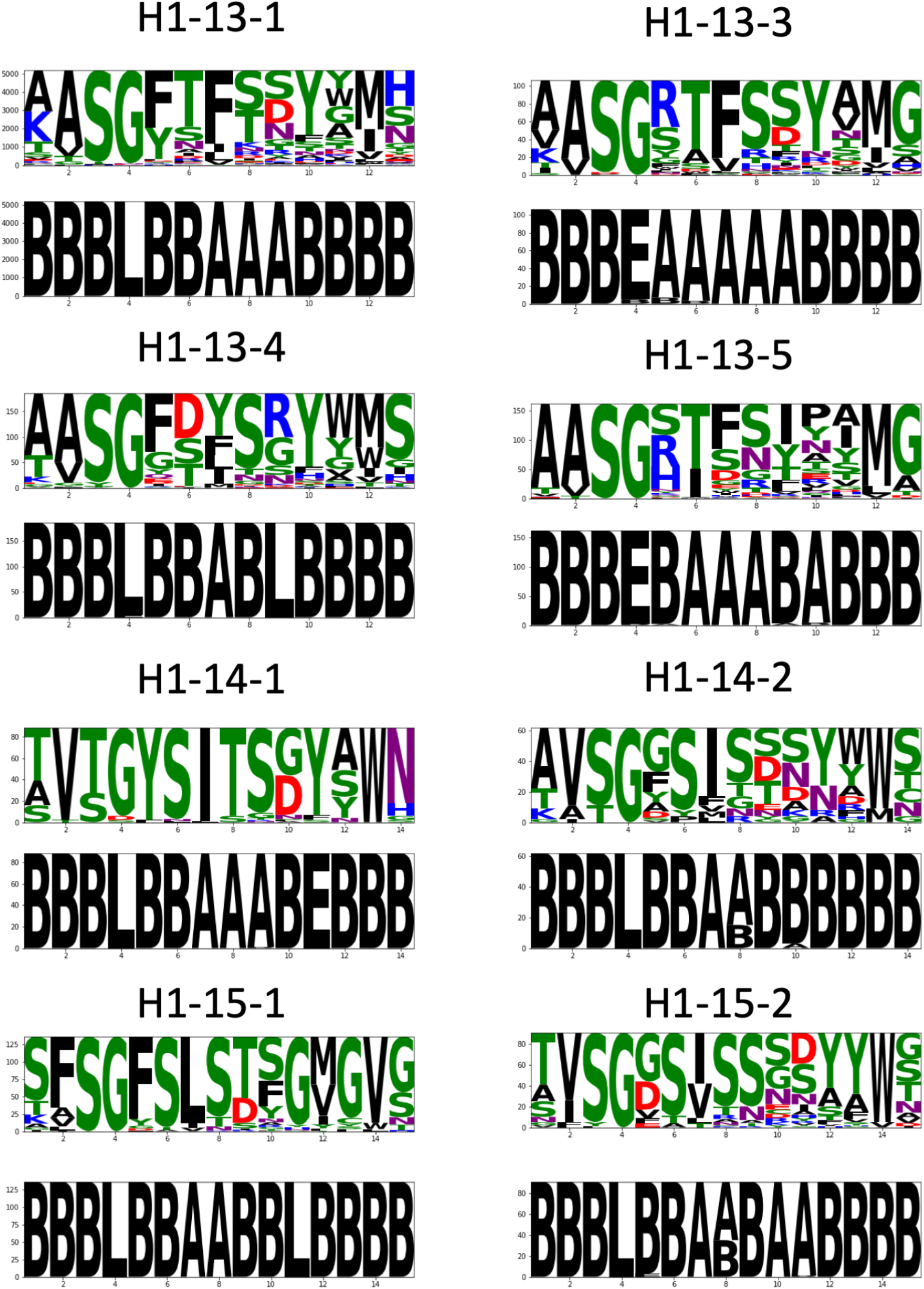
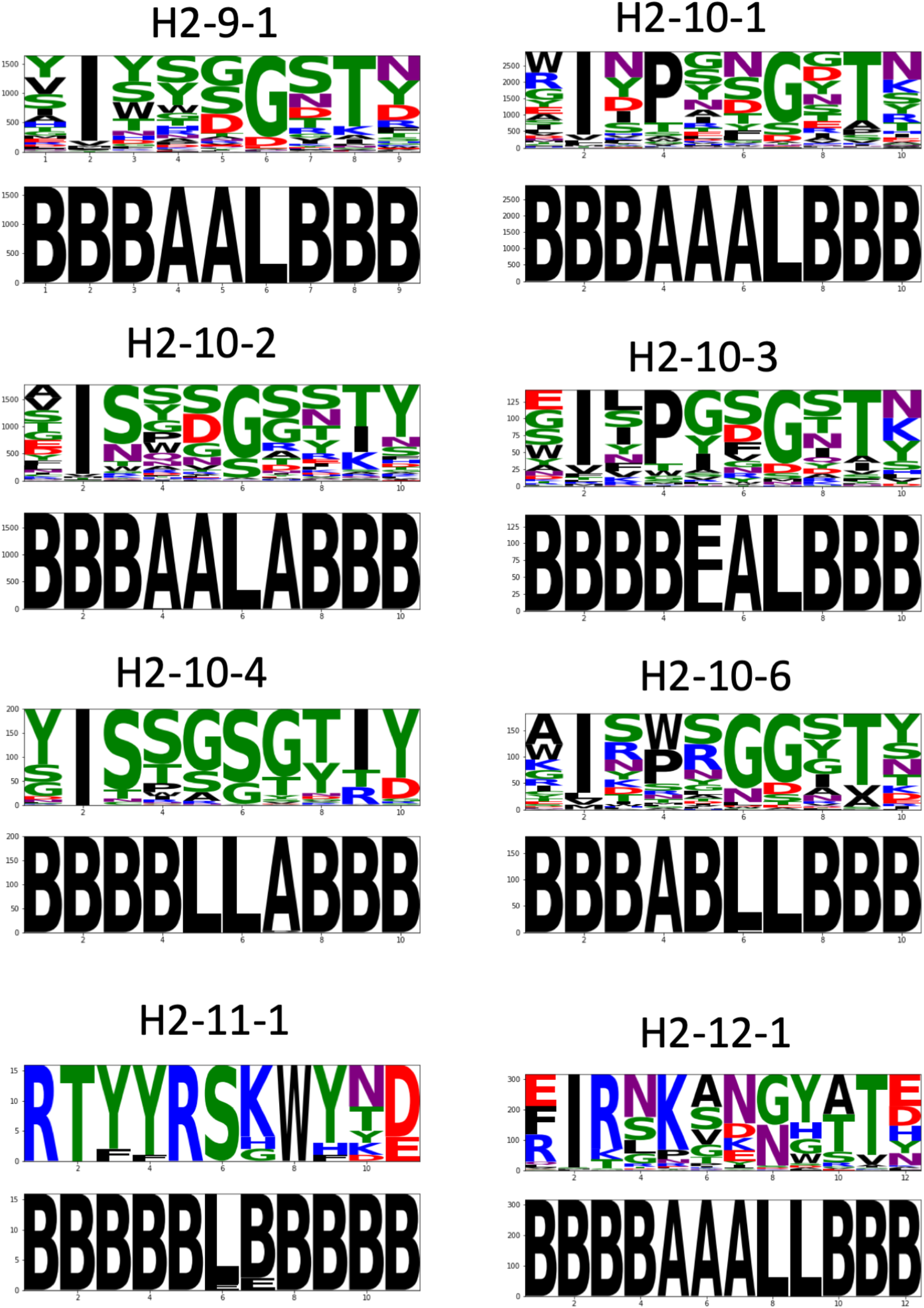
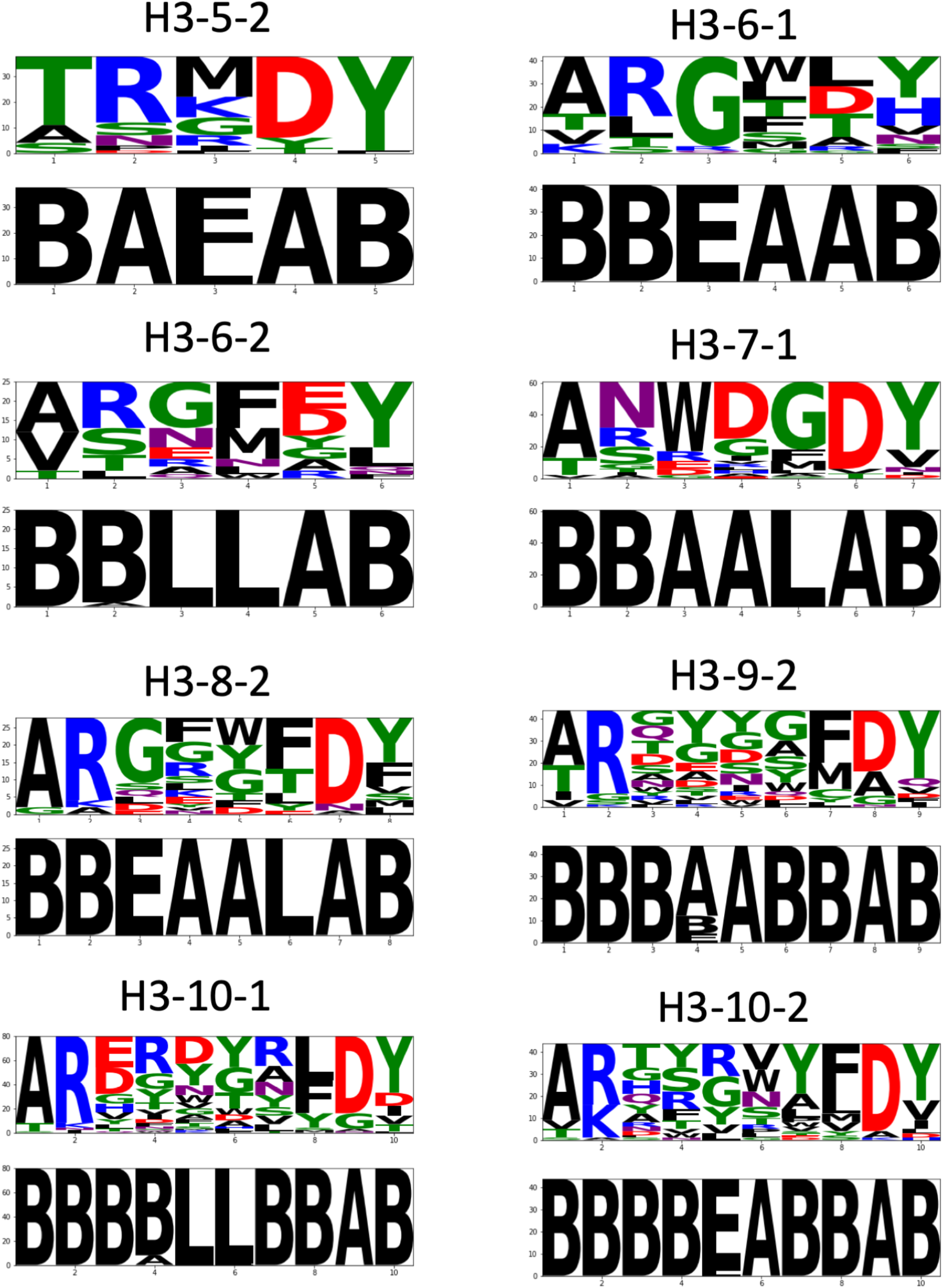
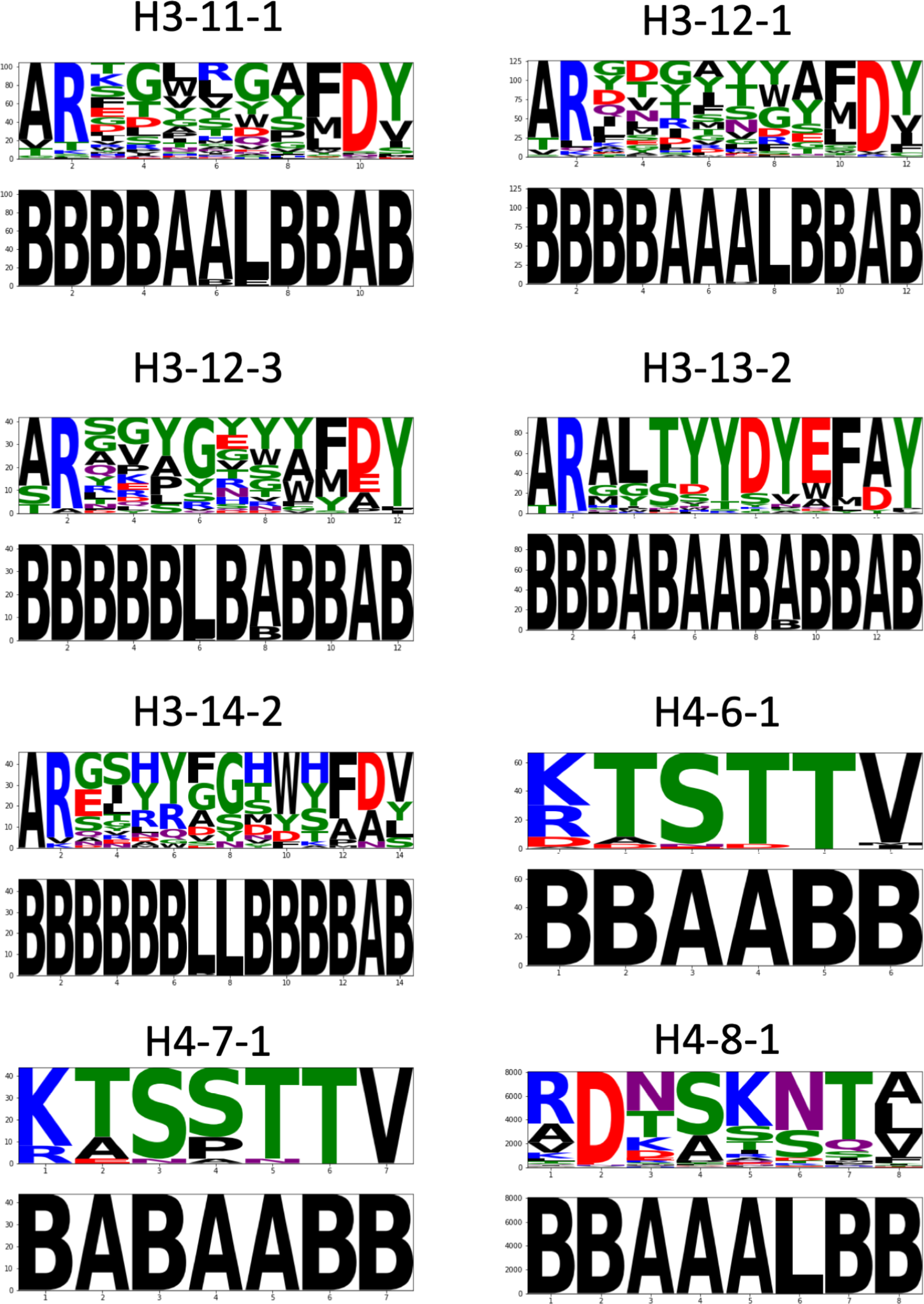
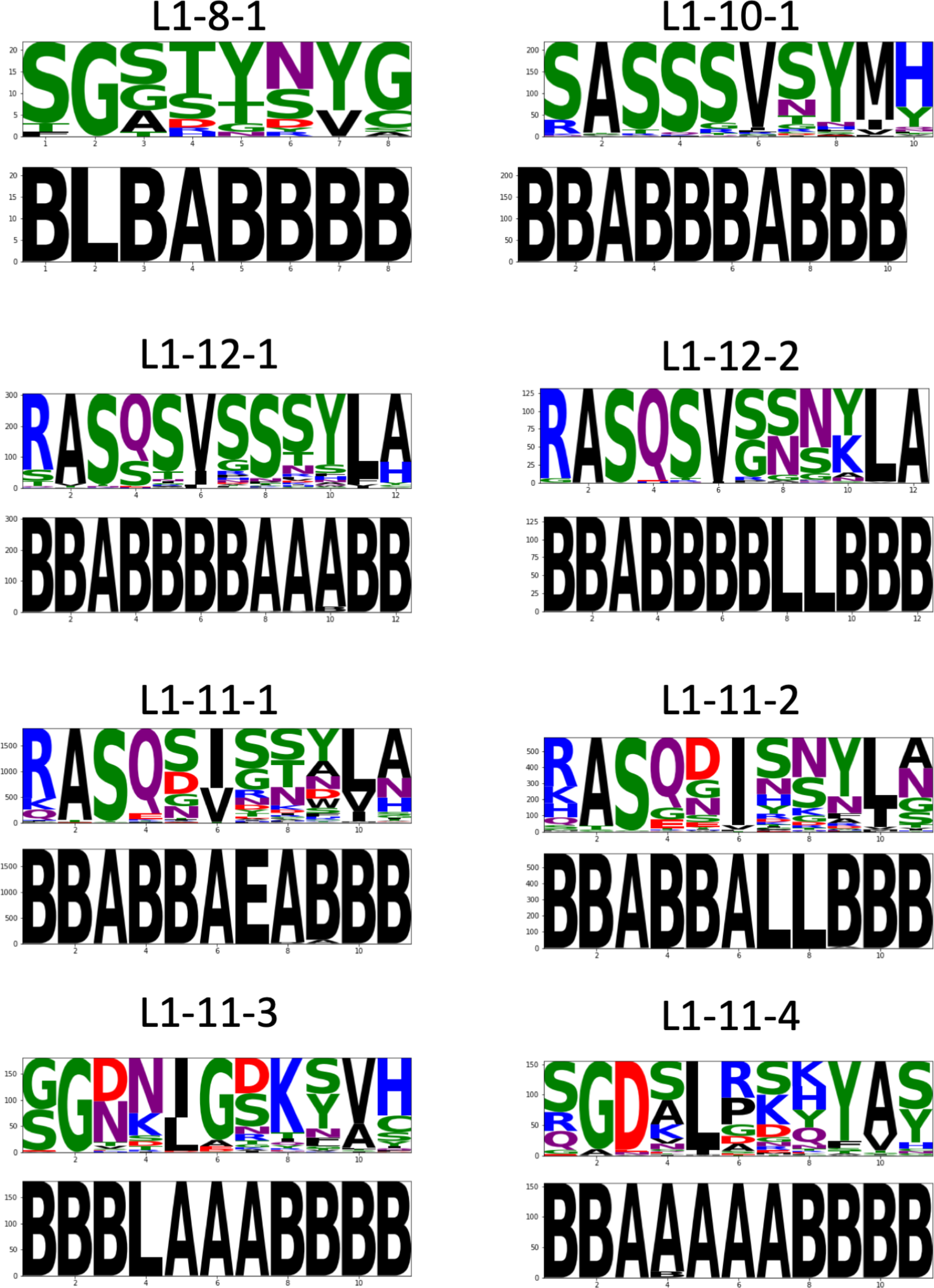
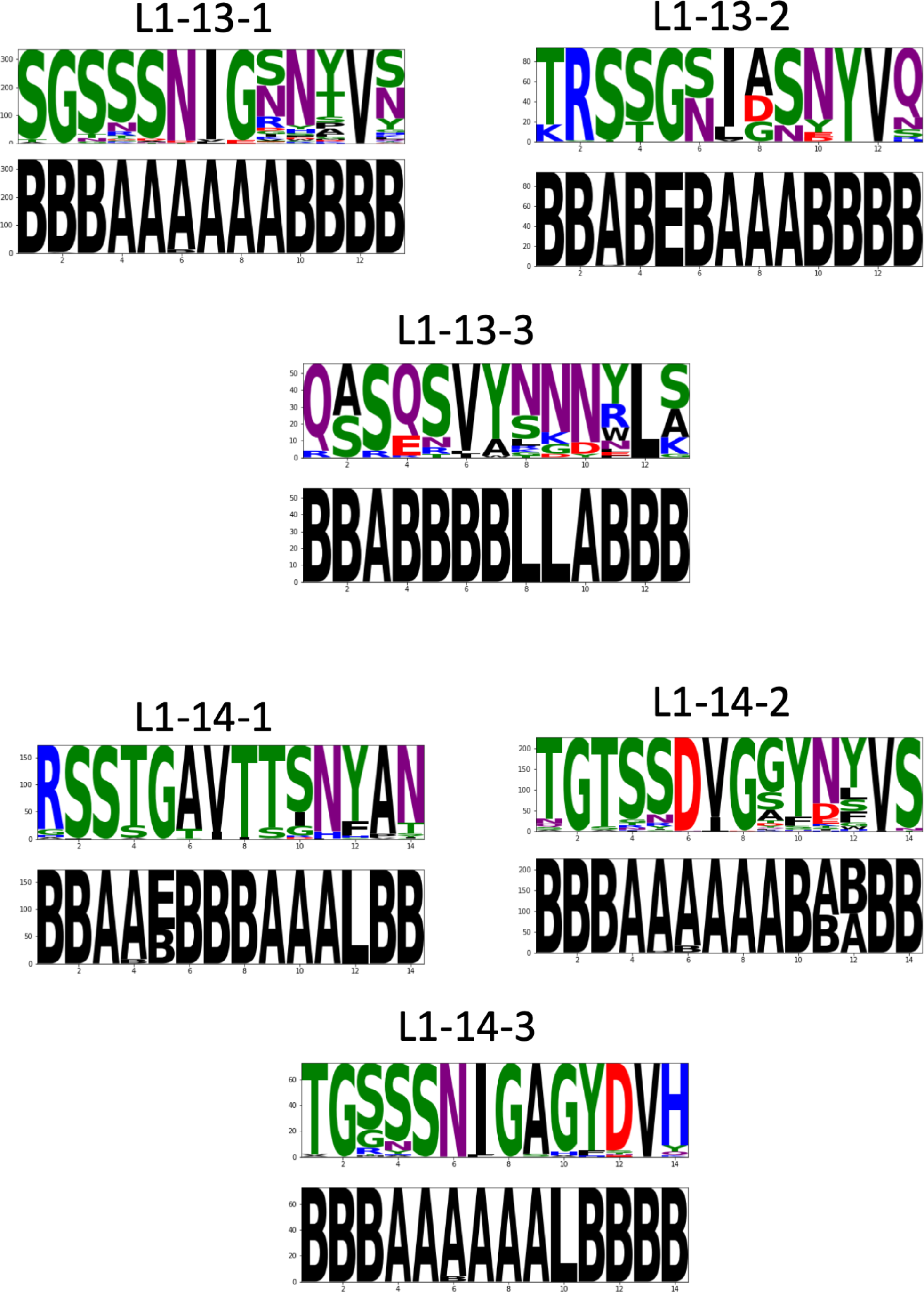
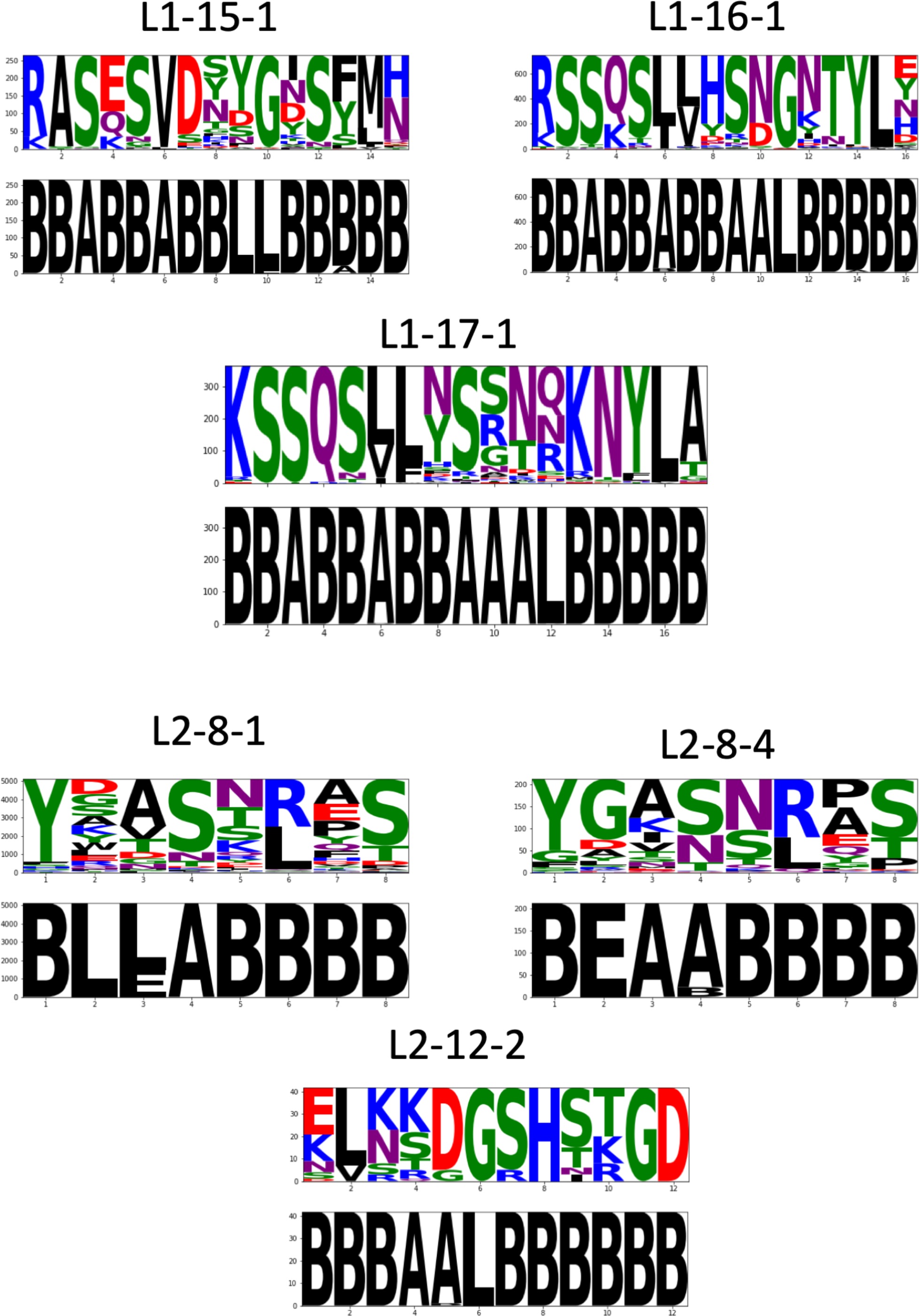
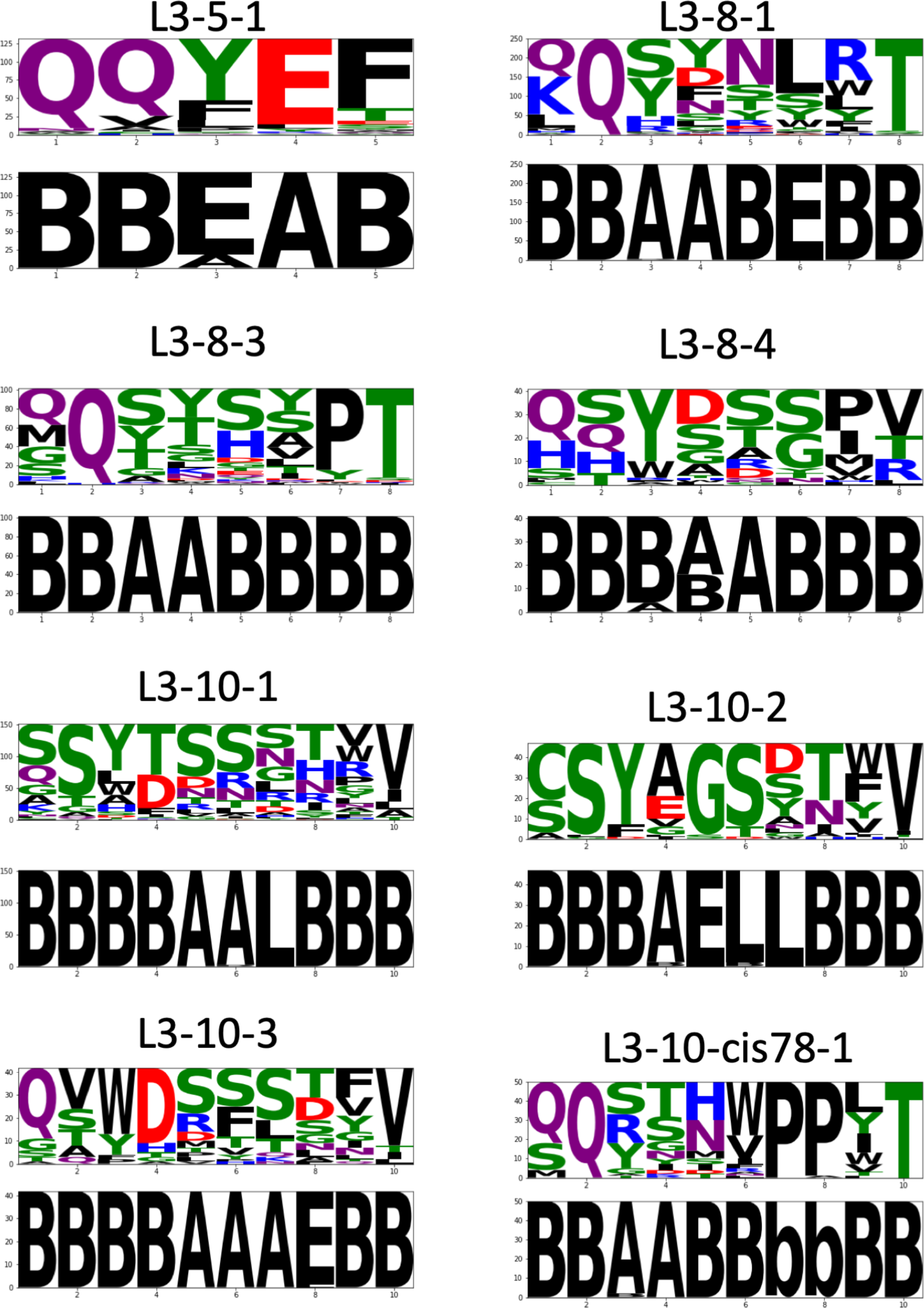
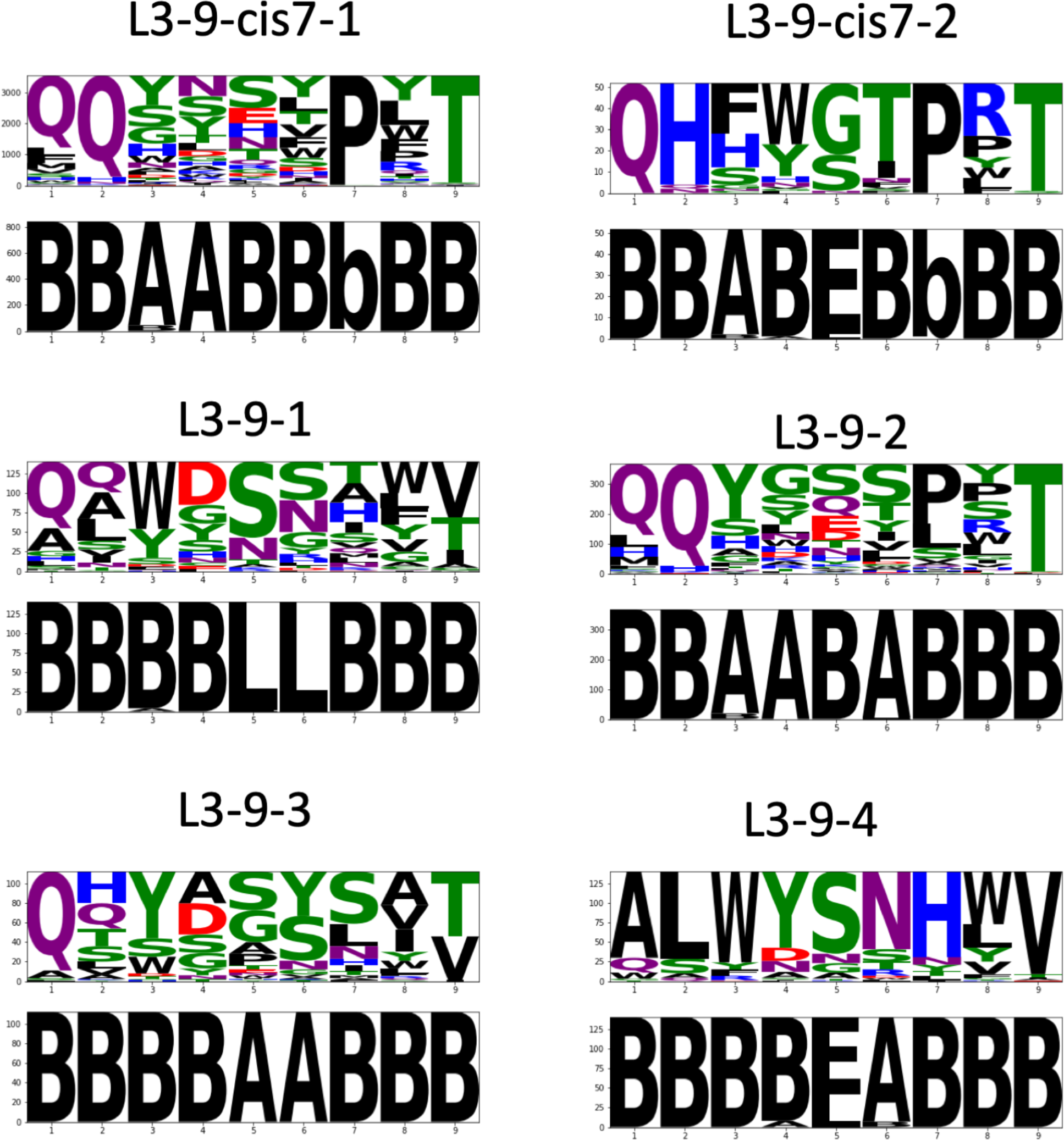
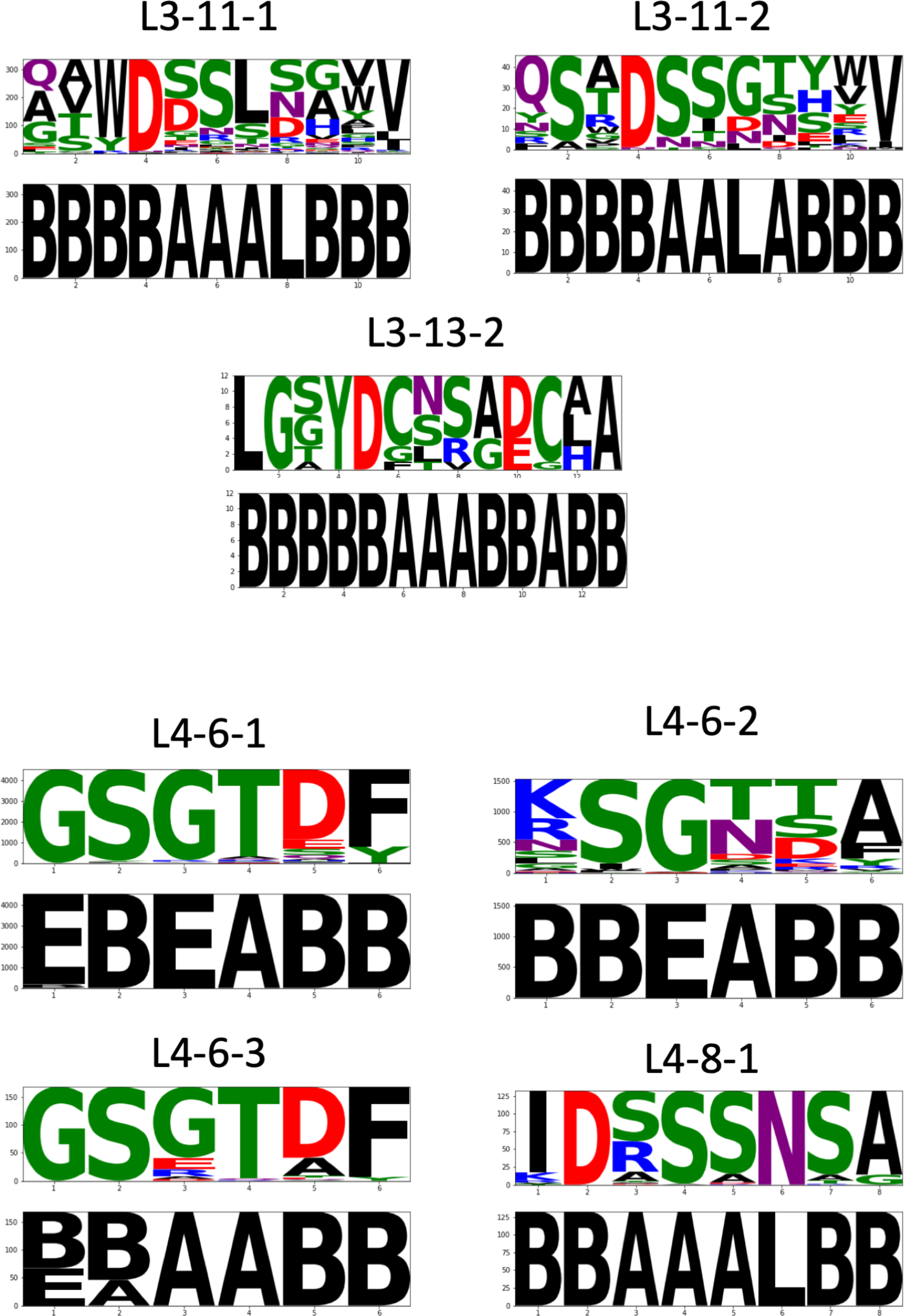
Sequence logos and Ramachandran region logos for the 2022 clusters. For the Ramachandran regions, “A” is the alpha helix region (ϕ<0°, −100° < ψ ≤ 50°); B is the beta sheet region (ϕ<0°, ψ>50° or ψ≤ −100°). L is the left-handed helical region (left-handed helices do not exist but the name has stuck: ϕ≥0°, −50° < ψ ≤ 100°). E is the “epsilon region” or (ϕ≥0°, ψ>100° or ψ≤ −50°).

**Figure 8.**
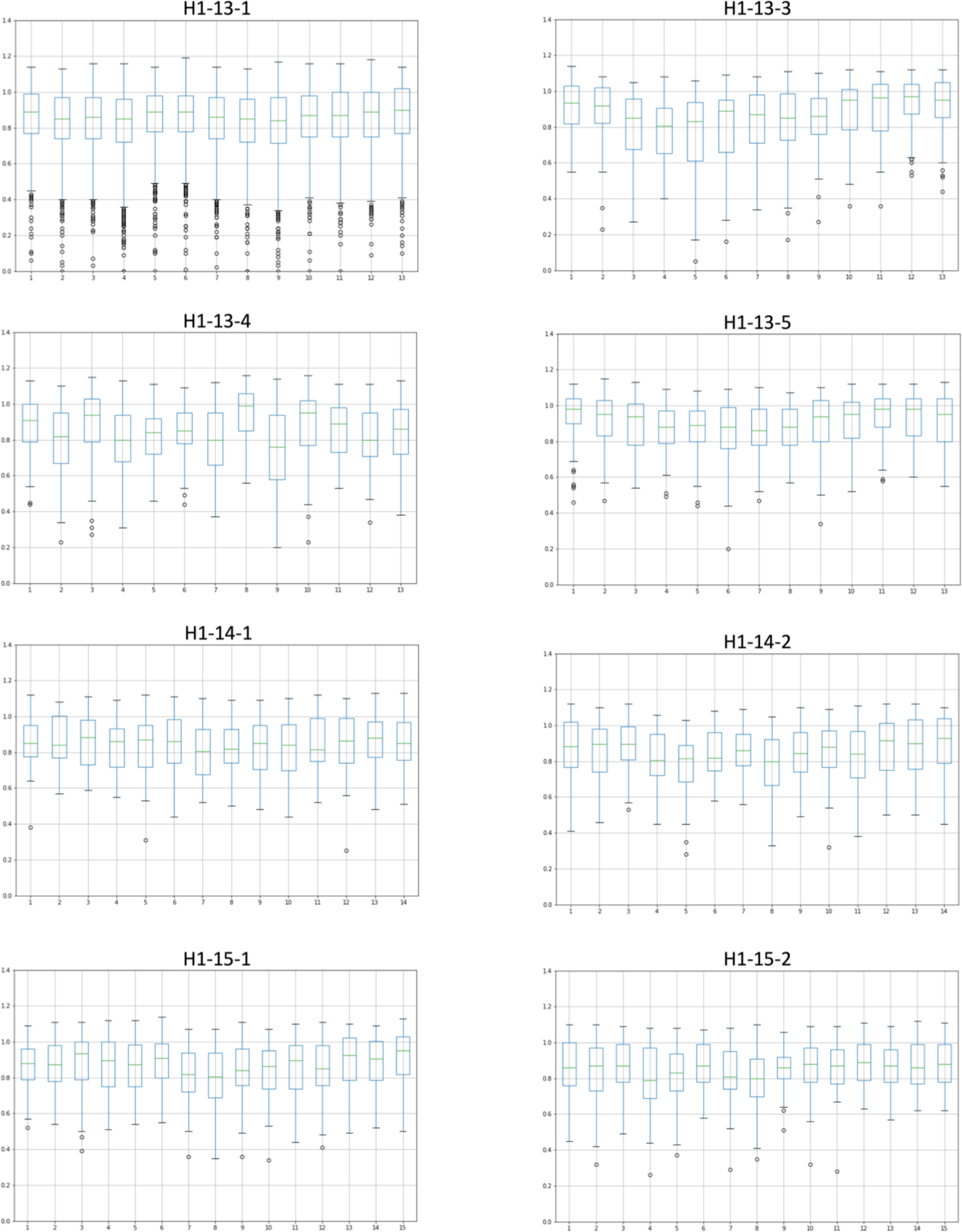
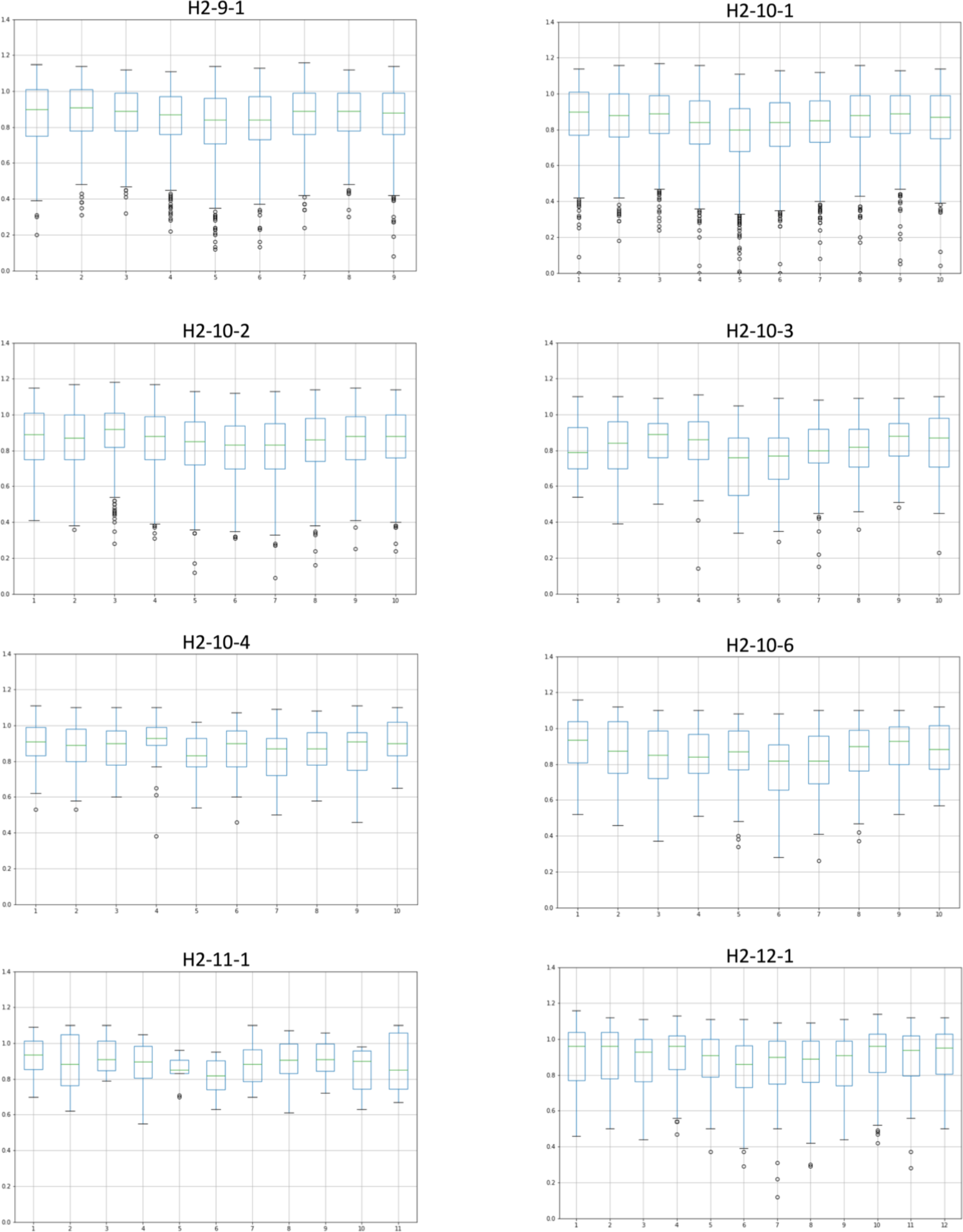
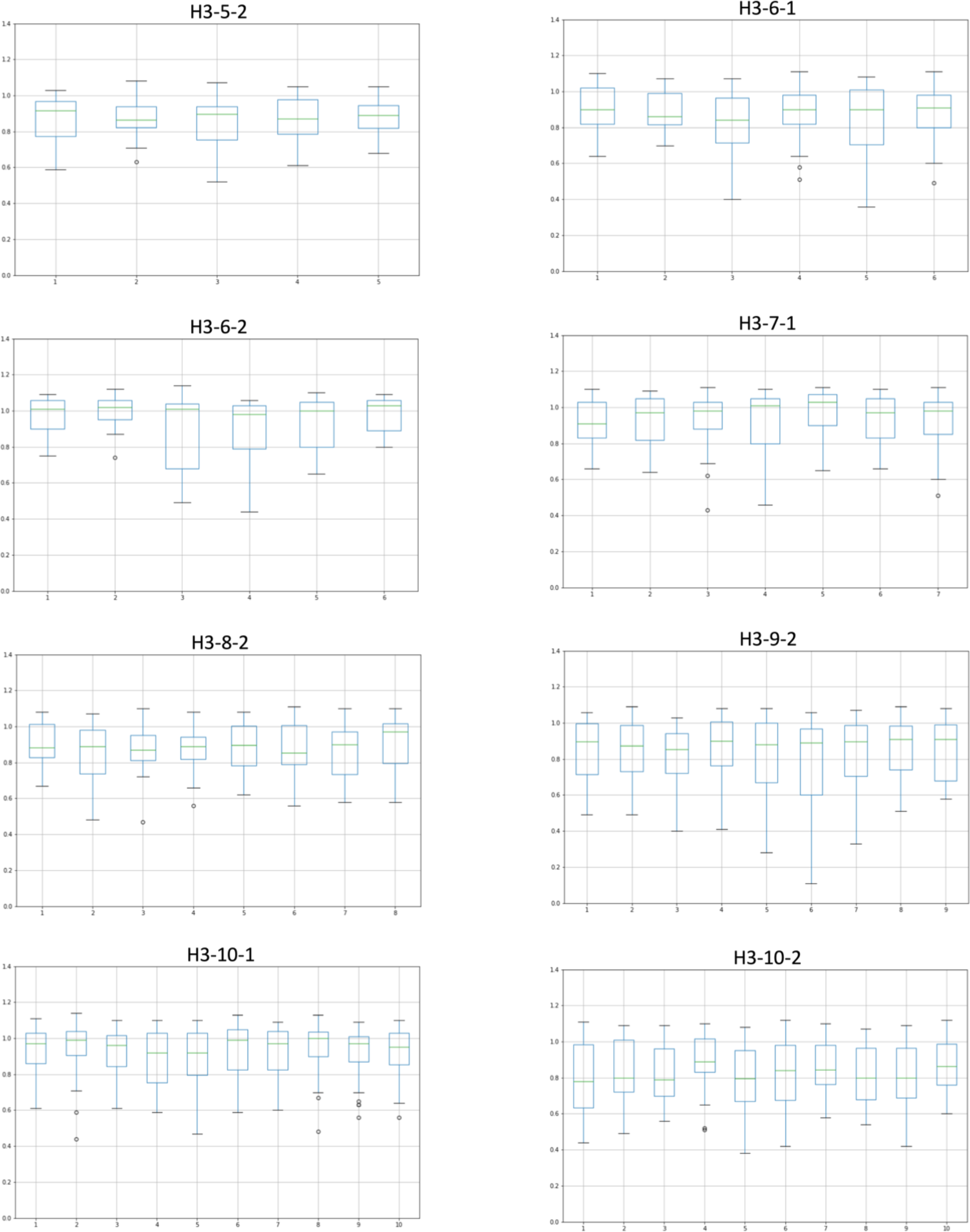
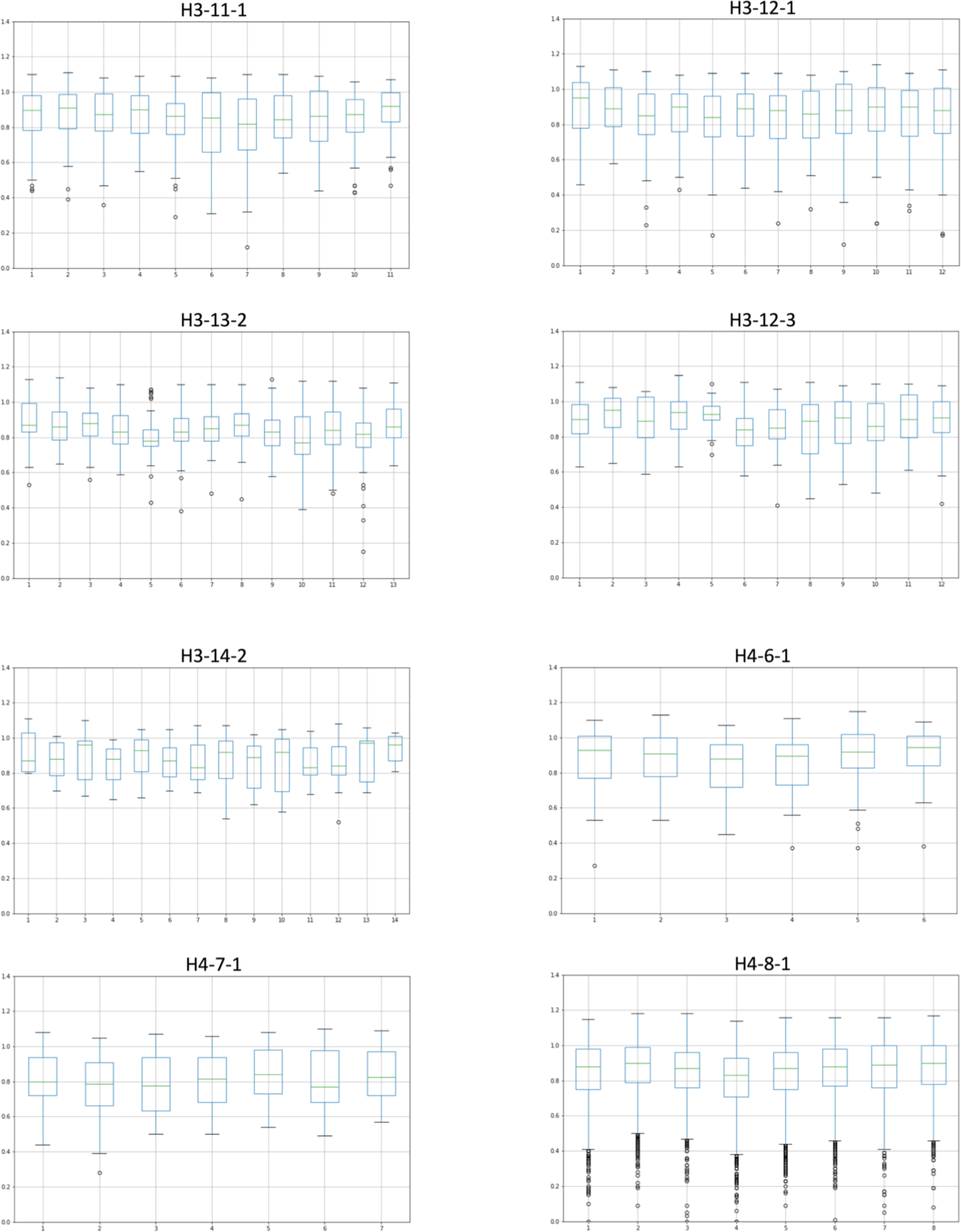
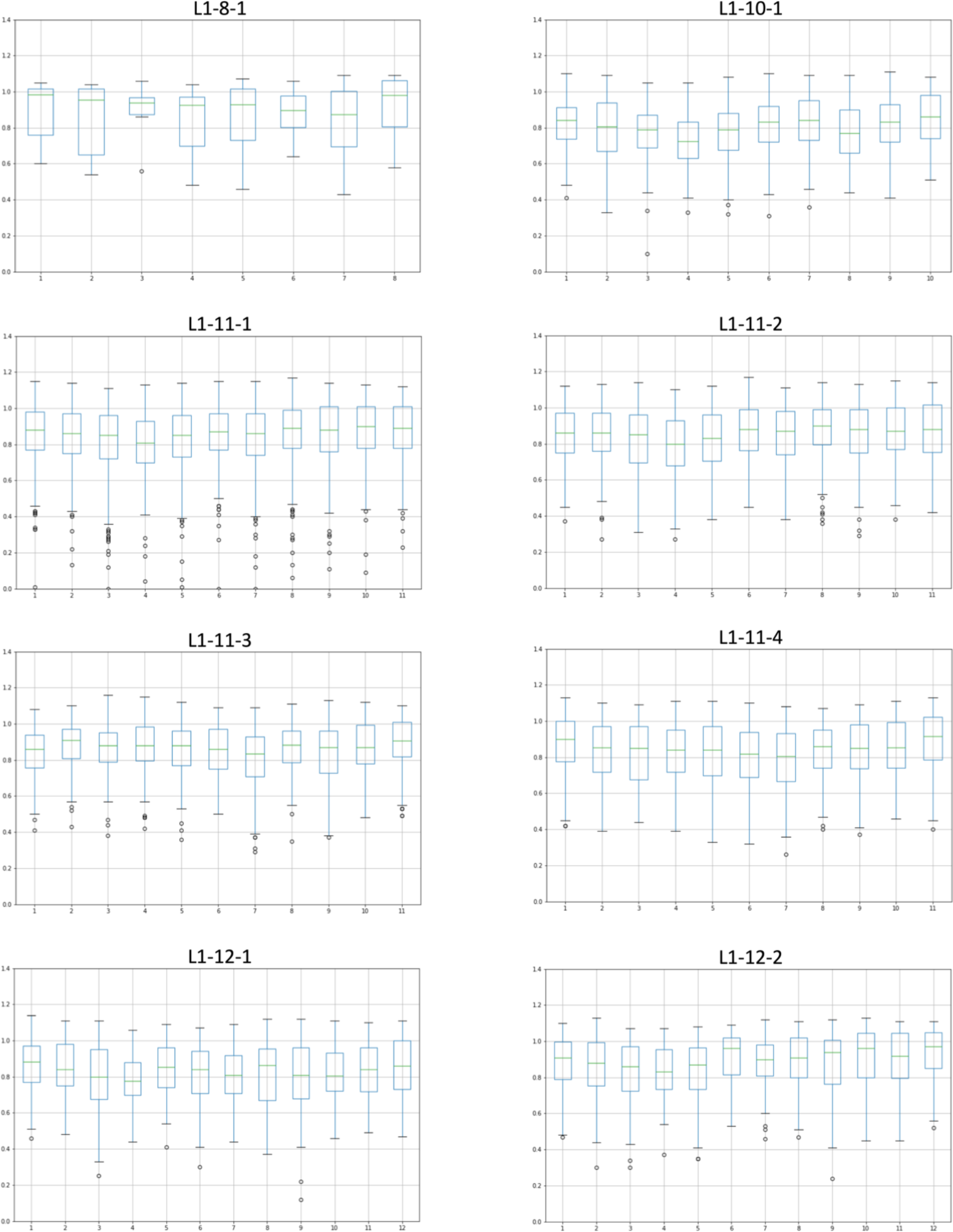
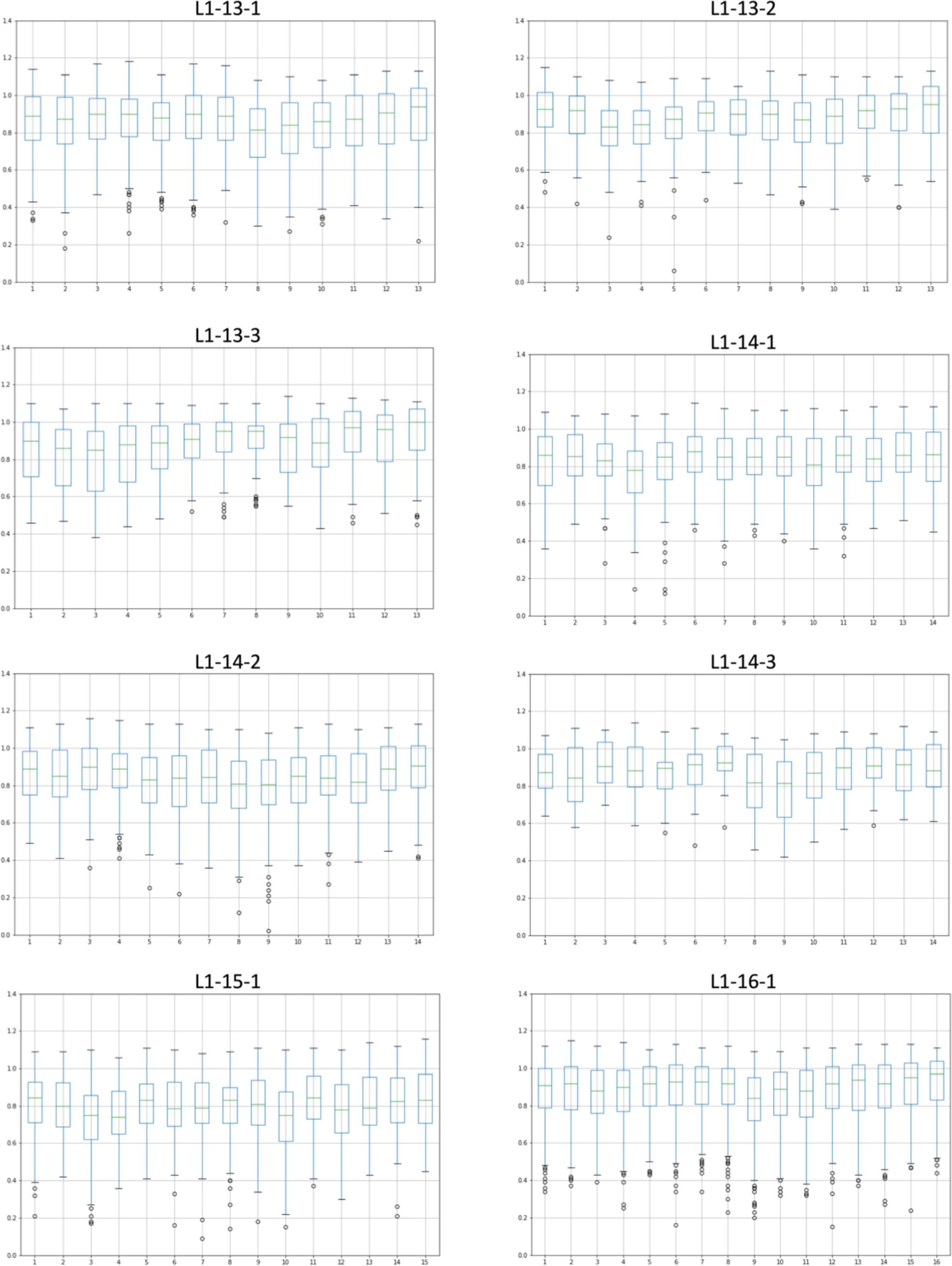
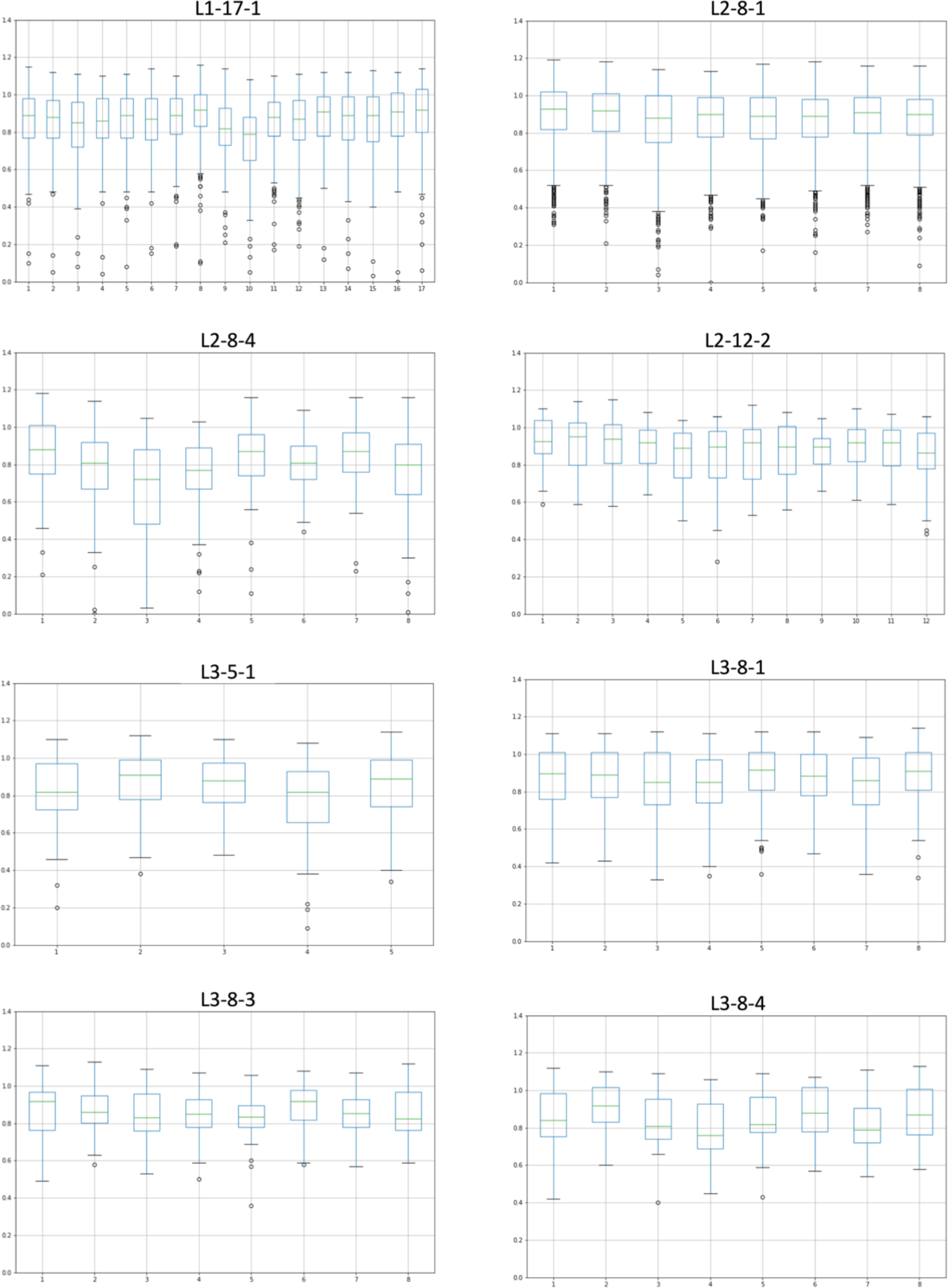
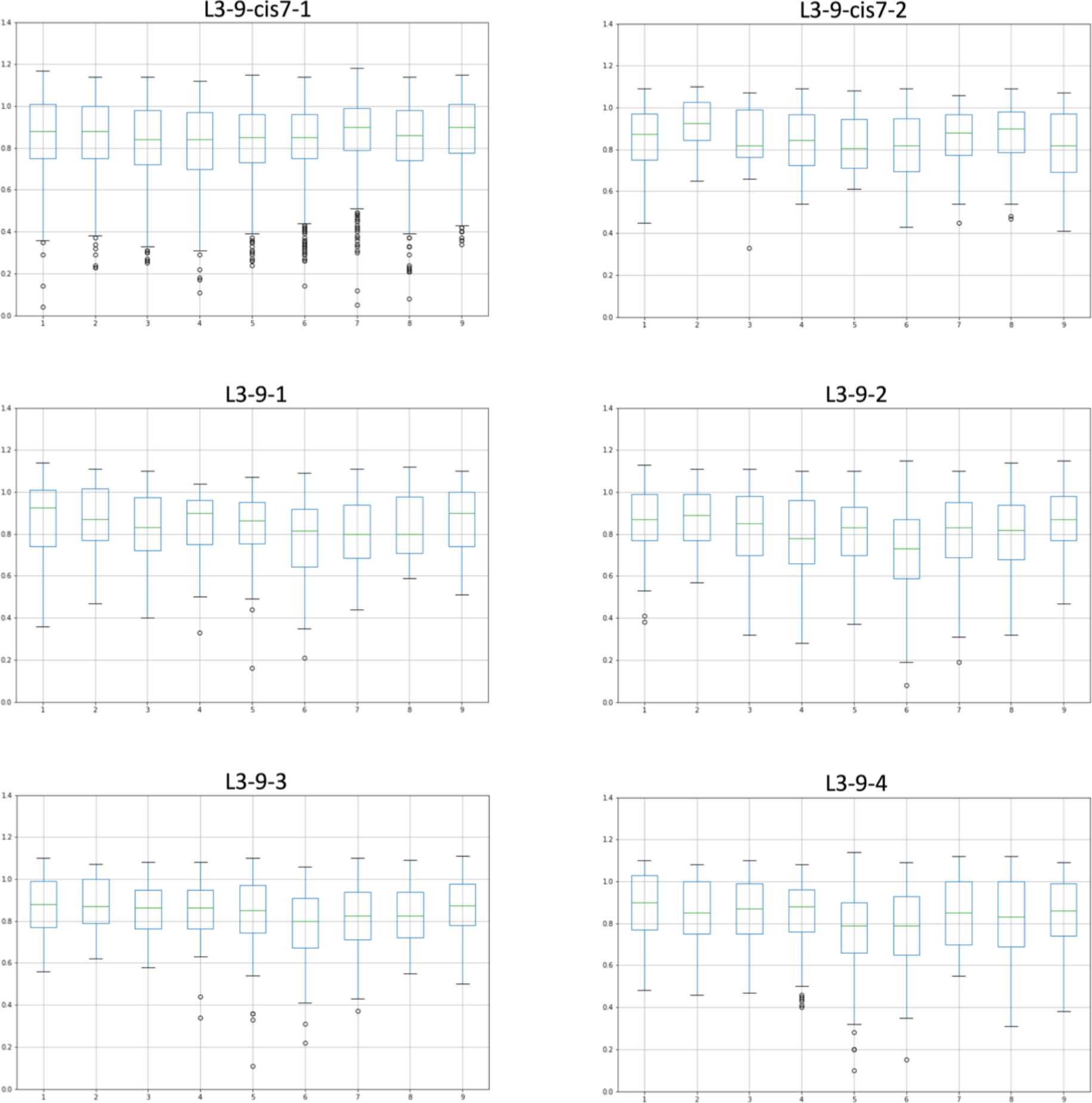
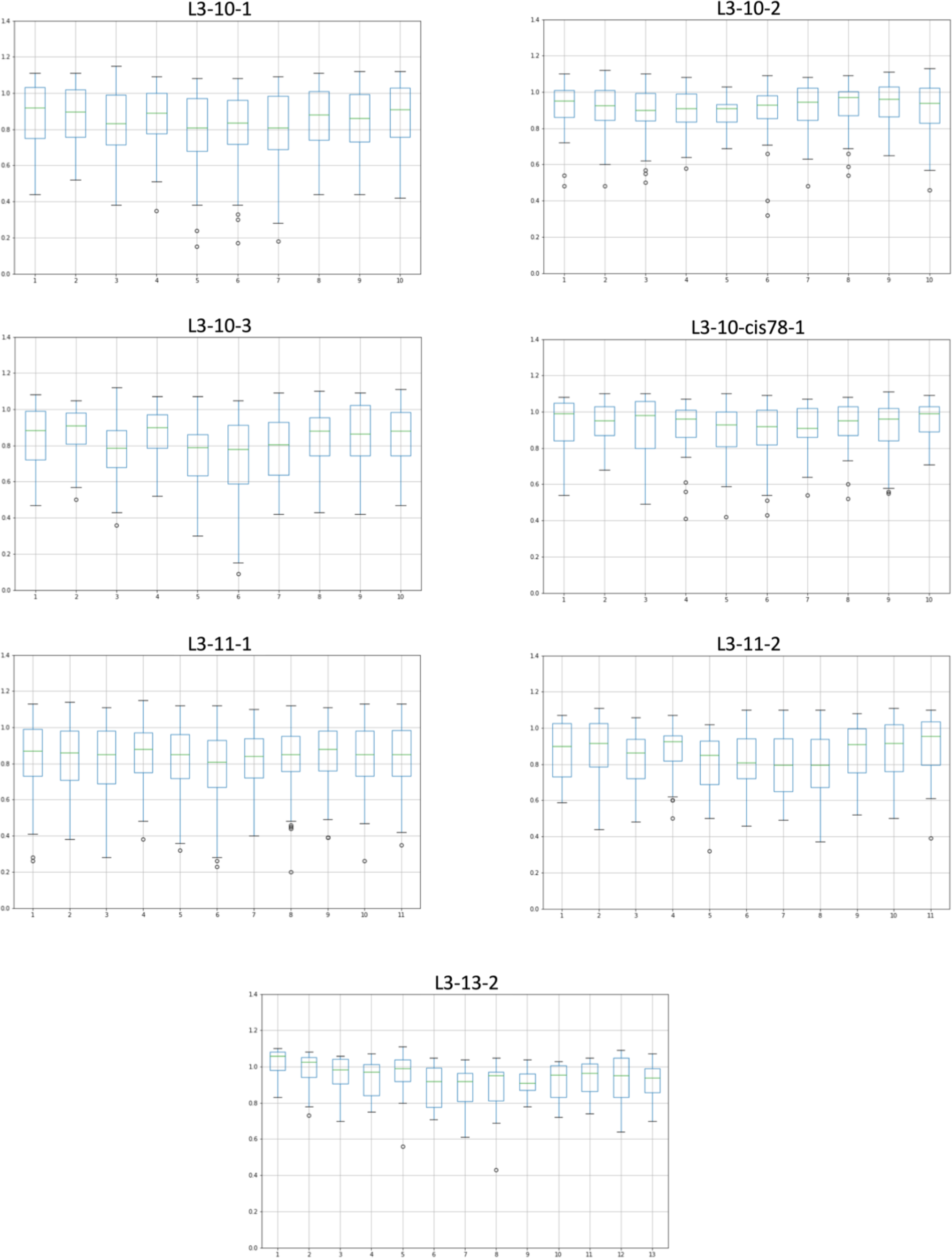
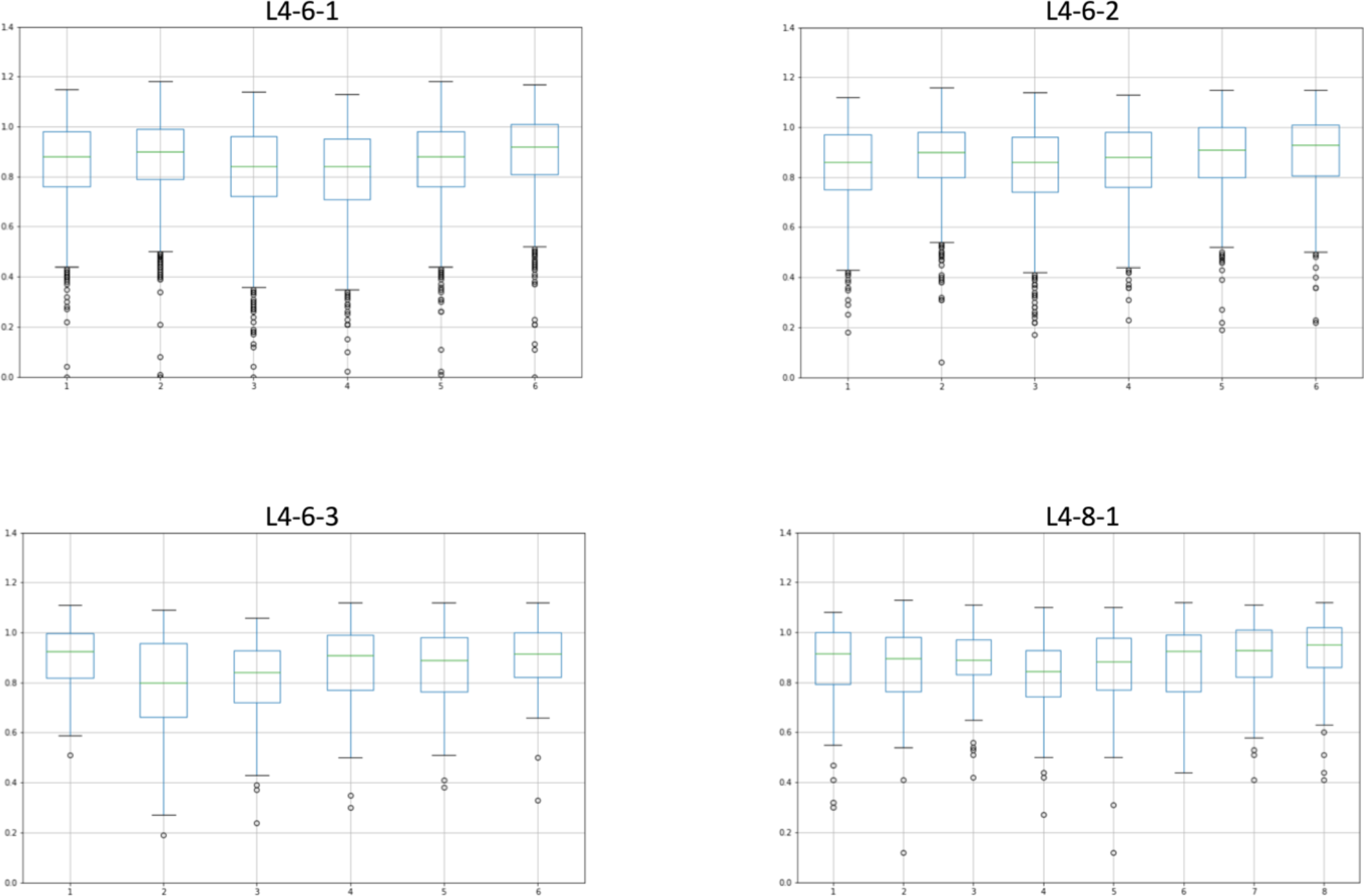
EDIA distributions for each cluster. Each y-axis is on the same scale.

Since the EDIA=0.0 clustering did not include NMR structures or X-ray or EM structures of resolution worse than 3.5 Å, we need a method to assign the remaining CDRs to our clusters. Such a method can be used to update our clusters periodically. To be consistent with density-based clustering, which is based on the number of neighbors of each point within a certain distance, we applied a nearest-neighbor approach (Benzécri 1982). We used the EDIA=0.0 clusters as “ground truth” and assigned each CDR to a cluster if it had a nearest neighbor in the ground truth set (but not from the same PDB entry) within a distance of 40° using the maximum dihedral angle metric. For large clusters, we limited the number of ground truth members to 1000 to speed the calculations. If no member of a ground truth cluster was within 40°, the CDR is assigned to noise (denoted with an asterisk, e.g. H1-13-*). Thus each CDR in a cluster has all of its backbone dihedral angles (ϕ,ψ,ω) less than 40° to a member of the ground truth set. As mentioned above, averaging dihedral angle differences can place CDRs with a peptide flip relative to the centroid if the cutoff for the average dihedral is too large (we previously used 40° for this cutoff). The maximum dihedral metric avoids this problem, and is a much stricter criterion than the average dihedral.

The resulting clusters are presented in Tables 5 and 6 for the CDRs of the heavy and light chains respectively. In total, there are 73 clusters across the 8 CDR loops. For the H1, H2, L1, L2, and L3, for which we had 72 clusters in 2011, we now have 52 clusters, of which 16 are new and 36 are the same as defined in North et al. The new clusters are denoted in blue type in Tables 5 and 6. The retired clusters are denoted in red in Table 1. The total numbers of clusters in the final set for each CDR are as follows: H1 (8); H2 (8); H3 (13); H4 (3); L1 (17); L2 (3); L3 (17); L4 (4).

**Table 5.**
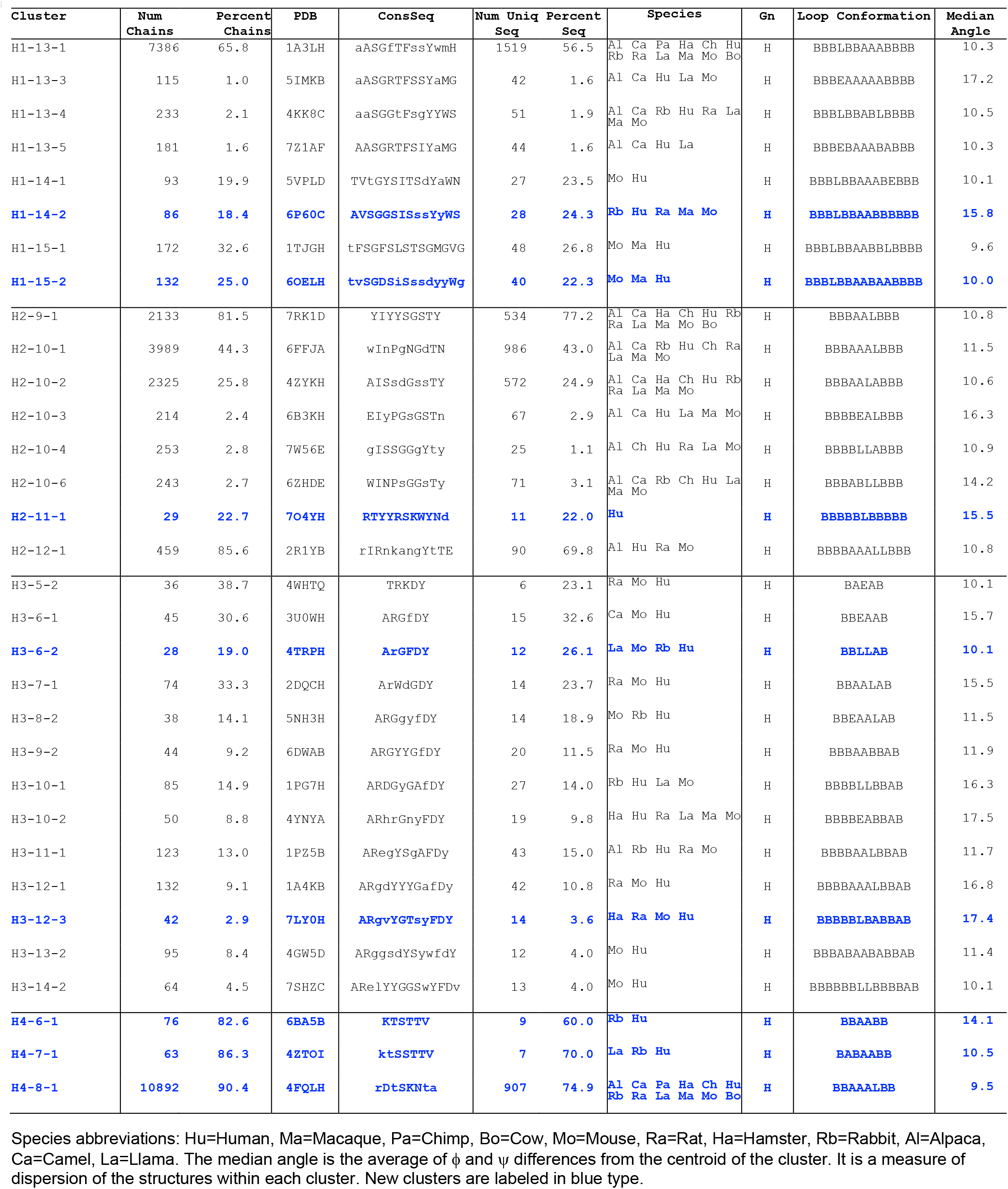
Final 2022 heavy chain clusters for PyIgClassify2.

**Table 6.**
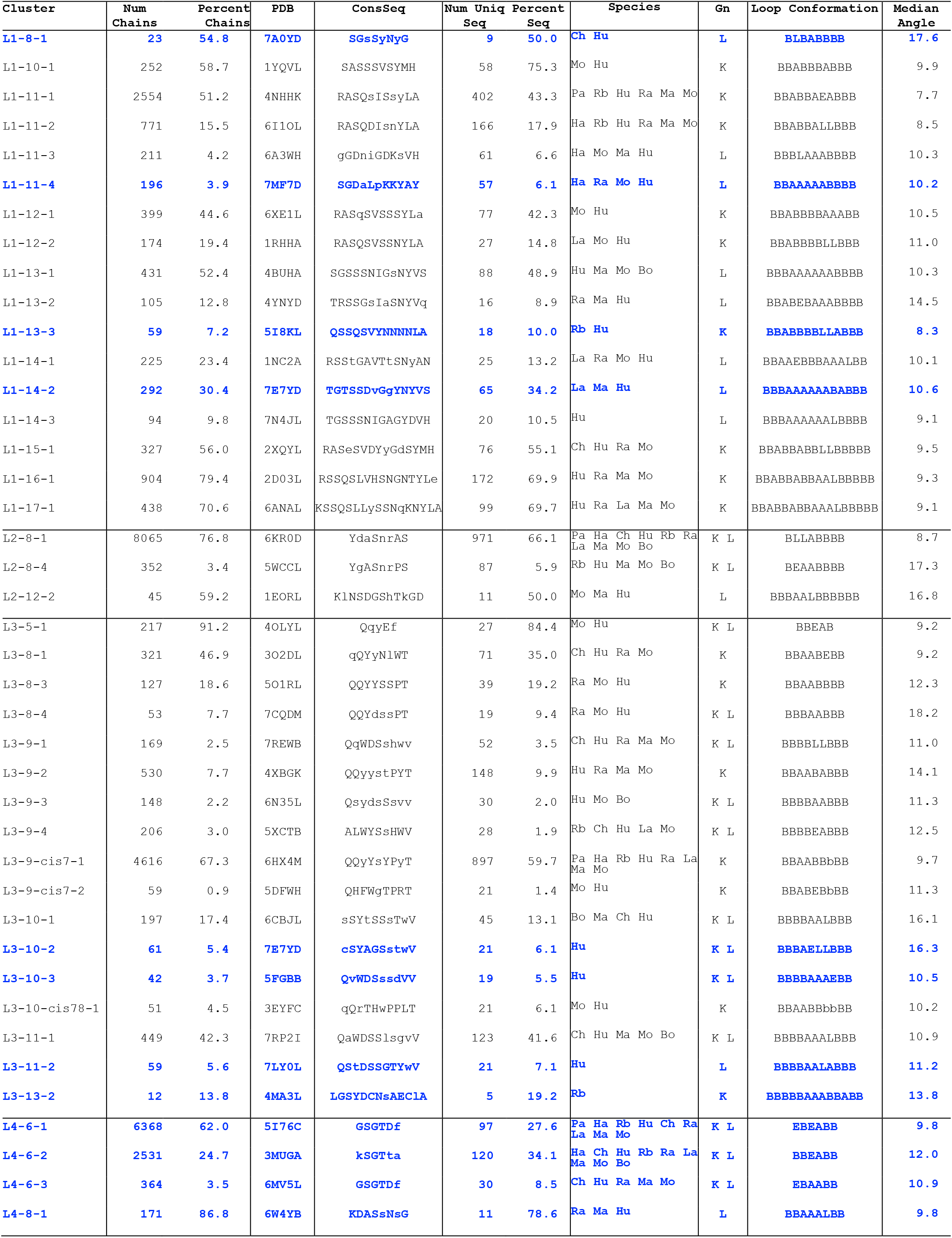
Final 2022 light-chain clusters for PyIgClassify2.

### Clusters related to each other by peptide flips

We identified all of the clusters that are related to each other by a peptide flip. Table 7 summarizes these clusters, listing which clusters are related, what their Ramachandran strings are, and what the peptide flip type is. In principle there are 8 different flip types: AA ⟷ BL; AB ⟷ BE; AL ⟷ BA; AE ⟷ BB; EA ⟷ LL; EB ⟷ LE; EL ⟷ LA; EE ⟷ LB, since in first position a change of ψ by 180° results in A⟷B or E⟷L and at second position a change of ϕ by 180° results in A ⟷ L or B ⟷ E. For example, L1-11-1 and L1-11-2 are related by an EA→LL flip, due to hydrogen bonding when the last residue of the L4 loop is Tyr (L1-11-2) instead of Phe (L1-11-1). This has been well known since the 1990s (Al-Lazikani et al. 1997). The two H1-15 clusters represent a peptide flip from BL→AA at the 10^th^ and 11^th^ residue position within H1. The L3-9, largest cluster, L3-9-cis7-1 is related to cluster L3-cis7-2 via an AB →BE flip at positions 4-5. Several examples are shown in Figure 9.

**Table 7.**
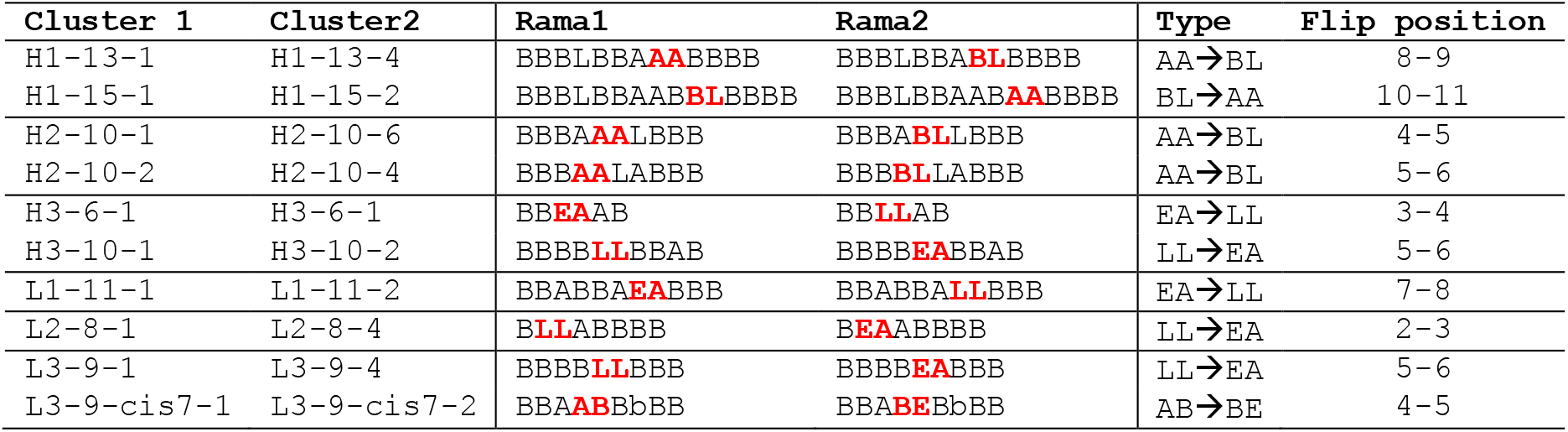
Flips between clusters.

**Figure 9.**
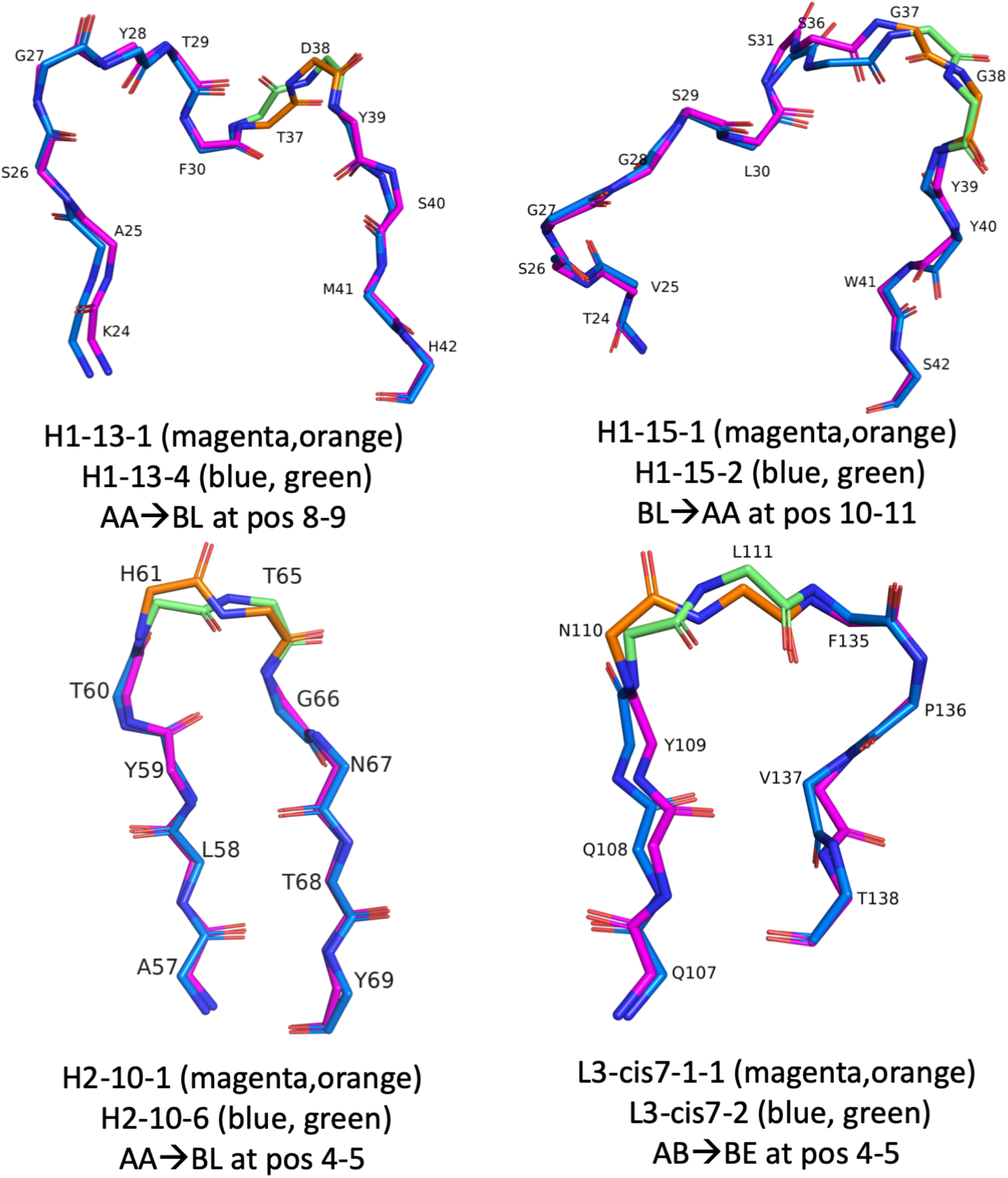
Cluster pairs with peptide flips (change in ψ by 180° at residue *N* and change in ϕ by 180° at residue *N*+1), without significant dihedral angle changes (>40°) at other positions.

### Updated website

The PyIgClassify website has been updated with the new data covering the PDB as of August 31, 2022. The site is located at http://dunbrack2.fccc.edu/PyIgClassify2. The download data are available under a CC-BY-NC license. Commercial users should contact the authors. The download data include:

1. File: `“pyig_cdr_data.txt”` which contains one line per CDR, including PDB information, cluster and distance values, sequence, Ramachandran string, germline assignment of the framework and the CDR (which may be different if the antibody is humanized) and their sequence identities, and the sequence of the CDR in the germline.
2. File: `“pyig_domain_data.txt”` which contains one line per variable domain, including the same information but for all four CDRs.
3. File: `“pyig_mmcif.tar.gz”`, a tar-gzipped file of mmCIF files for all variable domains in the PDB renumbered according to the modified AHo scheme.
4. File: `“pyig_cluster_mmcif.tar.gz”`, a tar-gzipped file of mmCIF files for all clusters separated into separate folders. Each file name includes the name of the cluster for ease of visualization in PyMol or Chimera (so that object names include the cluster identifier, for example: `H1-13-1_1H_2J88H_model1.cif`.
5. File: “pyig_vhvl_mmcif.tar.gz”, a tar-gzipped file of mmCIF files for all VH/VL domain pairs in the pDB renumbered according to our modified AHo scheme. In each file, the “author chain ID” is either H or L for the heavy and light chains respectively.

Software for determining cluster membership for input antibody structures will be made available in the near future. The current website allows the user to submit a structure for cluster determination using the average dihedral metric we used previously and a strict 20° cutoff (instead of the 40° we used in the original PyIgClassify website). A new website with enhanced functionality is in preparation.

## Discussion

Following selection of the final list of canonical clusters, we have now established a new classification of canonical conformations of antibody CDRs that is rigorously validated by electron density calculations and sufficient sequence representation in the PDB. Many clusters from North et al. are now obsolete, since they represent either too few sequences or poor electron density at specific residues indicating likely misfitting of electron density, usually in relation to the largest clusters.

We have performed assignments of IMGT germlines to the framework and CDR sequences for all antibodies in the PDB. These data are provided in the download files on the website, and may be used to establish relationships of the germline sequences and their common somatic mutations with their observed structures in antibodies. This analysis is complicated by the presence of somatic mutations and will be provided at a later date.

As deep learning approaches advance for both antibody structure, comparisons of predicted structures with experimental structures can be analyzed with regard to whether they reproduce the sequence-structure relationships observed in the canonical clusters of CDRs rigorously derived from PDB data. As computational antibody design methods mature, it will be useful to determine whether structures are being designed that mimic naturally encoded antibodies from the germline (or their somatic mutations) or whether new conformations are being designed and observed in the designed structures. With our new clustering, such inferences are put on a firmer statistical footing, which may help in further development of antibody design.

## Methods

### Sequences and structure files

We followed the methods described in Adolf-Bryfogle et al. (Adolf-Bryfogle et al. 2015) to identify antibody variable domains in the PDB and to renumber the files according to the modified Honegger-Plückthun numbering scheme we used previously. The AHo scheme differs from IMGT by adding two numbers to residues after CDR1. We developed new HMMs for the heavy-chain, lambda-light-chain, and kappa-light-chain variable domains from structures in the PDB to improve the accuracy of identification of CDR segments and framework regions. Some unusual CDRs or antibodies with framework insertions were misaligned with our previous HMMs. This occurred particularly for bovine antibodies, which have unusually long CDR H3s. In addition, we produced new HMMs for the alpha, beta, gamma, and delta chain variable domains of T-cell receptors to distinguish these domains from antibody light-chain domains. We searched PDB sequences (from out PISCES server, http://dunbrack.fccc.edu/pisces/download/pdbaa) (Wang and Dunbrack 2003, Wang and Dunbrack 2005) with all seven HMMs using hmmsearch (Eddy 2009). Most domains appear in more than one HMM output file. The largest score for each domain was identified among the HMM output files. If this score was over 80.0, then the domain was assigned to that type of variable domain. This cutoff appropriately distinguishes true antibody domains from other V-type immunoglobulin domains in the PDB (e.g., CD4, CD8). An additional HMM (labeled “P”) was developed for some light-chain sequences that were otherwise misaligned by the kappa HMM.

Variable domain coordinates were extracted from the mmCIF format files from the PDB, and renumbered according to the HMM alignment produced with hmmsearch. Each domain was placed in a separate file; all chains in all entries were processed in this way. The domains are labeled with a “datatag” consisting of the numbered domain, the HMM type, and then the PDB entry and chain ID. For example, the light and heavy chain domains from PDB entry 2J88 are labeled 1K_2J88L and 2H_2J88H respectively. To account for multiple models in NMR structures, the model numbers are attached to the data tags and filenames, e.g., 1K_2J88L_1 and 1K_2J88L_model1.cif, respectively. Any CDR structures that had breaks in the backbone polypeptide chain or missing coordinates were identified, and subsequently discarded from the post-clustering analysis.

The renumbered mmCIF files are available for download from the PyIgClassify website.

### Maximum dihedral angle metric

The work in North et al. did not use the RMSD calculation as a metric for comparing two loop conformations, but instead used a metric based on dihedral angles. Martin and Thornton previously clustered antibody CDRs based on dihedral angles (Martin and Thornton, 1996), but North et al. calculated the difference between two corresponding dihedral angles with a formula taken from the field of angular statistics that accounts for the periodicity of torsion angles (Mardia and Jupp 2000). Specifically, for each residue in two loops being compared, North et al. averaged the following angular distance metric to compare the dihedral angles of corresponding amino acids between two different CDR loops:

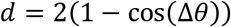

where θ is one of the protein backbone dihedral angles ϕ, φ. The data were presorted by their pattern of cis and trans residues, so that or ω did not need to be part of the average.

For clustering using DBSCAN, instead of the average dihedral metric, we use the maximum value of *d* between two different CDRs of equal length and take that as the clustering metric to compare two loops:

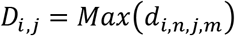

where *i* and *j* represent two loops of the same length being compared, and *n* and *m* represent corresponding dihedrals between those two loops. The maximum is taken over the ϕ, φ, and ω dihedral angles, so presorting by cis-trans pattern is unnecessary. The maximum dihedral metric is much stricter than the average dihedral metric, because it is sensitive to differences at a single residue, whereas averaging *d* over all residues will tend to balance out small differences at individual residues in favor of other residues being similar on average.

### Clustering with DBSCAN over a grid of its parameters

There are dozens of published clustering algorithms, some developed for very specific applications, while others are quite general and can be applied to a variety of scientific problems (Xu and Tian, 2015). The work in North et al. used the affinity propagation clustering algorithm (Wang et al., 2008), which was state-of-the-art at the time of publishing in 2011. This clustering algorithm defines clusters without needing to specify the number of clusters beforehand, but it does not account for noise points, which can distort the clusters. DBSCAN (Ester et al., 1996) is a density-based clustering algorithm that defines clusters by separating data points within high density separated by low density. A primary feature of DBSCAN is that it explicitly accounts for noise points, and collects them into a separate collection of data points. Noise points are points that do not lie anywhere near any of the defined clusters.

DBSCAN is a natural choice for this problem due to its robust ability to define well-resolved clusters, as well as the automatic detection of outlier data points, which are prevalent within the antibody CDR dataset. Outlier structures are mostly due to either unusual structures that have highly divergent sequence from rare germlines or synthetic antibodies, or errors in structure determination resulting in structures far from the canonical clusters.

With the selection of DBSCAN as the clustering algorithm, the selection of parameters for DBSCAN is key to generating a desirable set of clusters. DBSCAN requires two main parameters to run the algorithm, *MinPts* and ε. The algorithm is as follows:

1. Data points that have at least *MinPts* neighbors within a distance ε are labeled as *core points*.
2. Core points are connected by edges if they are within ε of each other.
3. Data points that are within at least ε of a core point are labeled border points, and an edge is placed between the border point and its closest core point.
4. Points which are not within ε of core points are labeled as noise.
5. The final cluster selections are the connected subgraphs of all of the core points and border points.

When first using DBSCAN with the angular distance metric defined in North et al., it was clear there were some selections of *MinPts* and ε resulted in a desirable clustering, while other selections either produced too much noise or resulted in clusters that were obvious merges of distinct conformations (e.g. distinct populations of different regions of the Ramachandran map for some residues). Some viable clusters with more diffuse density were observed at higher ε values, but these ε values sometimes merged clusters in undesirable ways. To address this issue, we developed an adaptation of DBSCAN that runs the algorithm over a grid of different values of *MinPts* and ε, and then combines the information over many different parameter sets to generate a final clustering. We call this adaptation Grid-DBSCAN (GDBSCAN). This is the method we used to cluster H4 and L4, and more information is provided in that paper (Kelow et al. 2020).

In order to combine results from multiple parameters selections for the DBSCAN algorithm, we implemented a graph theory approach. First, we treat each cluster output from a run of DBSCAN at a particular selection of MinPts and ε as a node on a graph. Second, we delete any clusters from each DBSCAN run that do not meet the criterion that the minimum distance between the furthest members of the cluster is below 150° at every dihedral. This technique removed clusters that merge different regions of the Ramachandran maps (e.g. A vs B, E vs L, A vs L, B vs E) whose centroids are roughly 180° apart in ϕ or ψ or both. The remaining nodes on the graph now represent all dense clusters from all runs of DBSCAN over the entire grid of MinPts and ε. Clusters arising from different parameters may of course be related. These nodes are then connected by calculating the overlap of their cluster memberships using the Simpson similarity index given by the following equation:

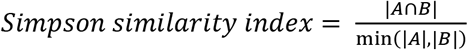

The Simpson similarity index will be 0 if there is no overlap between two clusters, and 1 if there is perfect overlap between two clusters or if one cluster is a perfect subset of a larger cluster. If the Simpson index is higher than 0.9, then an edge is drawn between these two clusters from different runs of DBSCAN. The final set of clusters is the set of all connected subgraphs within this larger graph structure.

Backbone clustering of the antibody CDRs was done using the following procedure. For each CDR-length, every CDR structure was included in the clustering set, and the grid for DBSCAN was set to 0.1 to 1.0, in steps of 0.1 for the parameter ε, and 5 to 20 in steps of 1 for MinPts. Some CDR lengths required a higher range of MinPts (up to 50 or 100), because of the large number of points available (L2-8, H4-8, L4-6).

### Electron density fit for individual atoms to support backbone clustering

Since the North clustering, strategies for handling quality assessment of protein structures in the PDB have become more robust (Fährrolfes et al. 2017, Meyder et al. 2017, Liebschner et al. 2019). Electron density support for individual atoms, or EDIA, was introduced in 2017 (Meyder et al. 2017). EDIA calculates the fit of the coordinates of individual atoms to their electron density in a spherical region around the atom and accounts for both positive and negative density in the electron density difference maps. Using EDIA presents an alternative to B-factor cutoffs or resolutions cutoffs, which are traditional data used to assess structure quality, and typically applied as a filter to cut out structures that do not meet a specific threshold. One primary concern with a resolution cutoff is that resolution is a value that summarizes the quality of the entire structure, but has no information on the scale of amino acid residues or atoms. On the other hand, B-factor does assess structure quality at the atomic level, but it is highly susceptible to errors during the structure determination and structure refinement step, and oftentimes correlates with non-quality related factors such as protein dynamics and crystal contacts (Schlessinger et al. 2006, Shapovalov and Dunbrack 2007, Yang et al. 2016). This makes the use of B-factor in the evaluation of protein crystal structure quality a less attractive option compared to EDIA. The EDIA data were downloaded from the ProteinsPlus webserver (Fährrolfes et al. 2017).

### Peptide flips from mis-solved residues within protein structures

Within segments of protein without regular secondary structure, a common phenomenon is an event called peptide plane flipping, where the backbone atoms of the protein backbone peptide plane configure in such a way that the carbonyl oxygen and backbone nitrogen are flipped 180° relative to a non-flipped counterpart (Hayward 2001, Touw et al. 2015). Peptide plane flipping is thought to have a role in protein conformational dynamics, but also are prevalent features in mis-solved residues. This may occur at low resolution when the two states may be difficult to distinguish, or due to molecular replacement with an incorrectly modeled structure. Peptide flips are determined by the co-dependency of the Ramachandran conformations of adjacent residues, where the ψ conformation of residue *i* is highly dependent on the ϕ conformation of the *i*+*1* residue. Given the importance of the backbone conformation in backbone clustering analysis, the opportunity to systematically identify features of mis-solved protein structures on the basis of EDIA calculations for each CDR atom, and the sensitivity of the DBSCAN clustering protocol to pick up minute differences at single dihedral resolution, we applied an analysis of protein backbone flips to structures within the CDR set to identify clusters that are related to each other by peptide flips, and relating the peptide flip feature to errors in protein structure determination.

To identify clusters that are related by peptide flips of the protein backbone, we started by calculating the average dihedral angle for each residue within each cluster of CDR structures. we calculate the average dihedral using the following equation for averaging torsional angles, which takes into account periodicity at 360°:

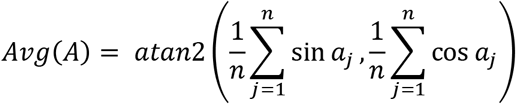

Following the calculation of the average dihedral at each residue within the CDR, we identify clusters that are related by peptide flips by considering sets of two neighboring residues within the CDR sequentially at corresponding residues between two loops. First we check that the ϕ and ψ angles of each residue before and after the residue in question are within 40° of each other in order to ensure that the motions observation between the two residues are related to peptide flipping and not natural backbone motion. Next, for the two residues being compared between two loops, we check to see if the difference in ψ of residue *i* within loop *j* is greater than 100°, and the difference in ϕ of residue *i*+1 is within loop *k* is greater than 100°. If all of the aforementioned criteria are met, position *i* is labeled as a flipped positions between the two loops, and the Ramachandran type for each of the two residues is labeled to capture the flip type. Flips between two clusters identified in this way were observed in PyMol and some pairs were in conformations that are more distinct than peptide flips and were skipped.

## Acknowledgments

This research was funded by NIH grant R35 GM122517 (to R.L.D.) and P30 CA006927 (to Fox Chase Cancer Center).

